# Dissecting Annexin-A11 into its functional domains revealed calcium as a key regulator for RNA transport and its association with ALS

**DOI:** 10.1101/2025.10.27.684738

**Authors:** Giulia Di Napoli, Lorenzo Alfurno, Alex Fissore, Eleonora Raccuia, Valeria Bincoletto, Paolo Olivieri, Riccardo Scoccia, Fabrizio dal Piaz, Mauro Marengo, Simonetta Oliaro-Bosso, Silvia Arpicco, Maela Manzoli, Gianluca Catucci, Gianfranco Gilardi, Adrian Velazquez-Campoy, Filippo Prischi, Angela De Simone, Francesco Di Palma, Francesca Spyrakis, Salvatore Adinolfi

**Author notes:** indicates first authorship (Giulia Di Napoli, Lorenzo Alfurno, Alex Fissore). denotes three co-corresponding authors (Francesco Di Palma, Francesca Spyrakis, Salvatore Adinolfi). **This work was supported**, by University of Turin (Ricerca Locale Grants ADIS_RILO_21_01, ADIS_RILO_23_01, ADIS_AUTOF_25_01, OLIS_RILO_22_02, MARM_RILO_22_05, MARM_RILO_20_04. SPYF_RILO_24_01, DESA_RILO_24_01. Programma di ricerca CN00000013 ’’National Centre for HPC, Big Data and Quantum Computing”, finanziato dal Decreto Direttoriale di concessione del finanziamento n1031 del 17.06.2022 a valere sulle risorse delm PNRR MUR-M4C2-Investimento 1,4-Avviso “Centri Nazionali” - D.D. n3138 del 16 dicembre 2021. Computing time was provided by the Data Science and Computation Facility at IIT on the Franklin HPC system under the umbrella of the Framework Agreement of June 26, 2023, between Fondazione Istituto Italiano di Tecnologia and University of Turin.

## Abstract

Recent studies reveal a “hitchhiking” mechanism in neurons, where organelles transported along microtubules carry other cargos via tethering molecules. Annexin A11 (ANXA11), a calcium-dependent phospholipid-binding protein, functions as a tether linking RNA granules to lysosomes, aiding mRNA transport for rapid neuronal responses. Structurally, its N-terminal (Nt) binds RNA, while the C-terminal (Ct) associates with lysosomal membranes. Mutations in ANXA11 linked to Amyotrophic lateral sclerosis (ALS) may disrupt this function.

Here, applying a multidisciplinary approach, we revealed that Ca^2+^ acts as a master regulator of ANXAll’s physiological function by modulating its conformational states. Specifically, Ca2^+^ influences a switch between two conformations: a close state, in which the Nt and Ct interact with each other, and an open state, which occurs in the presence of Ca2^+^ ions, where this self-interaction is disrupted, allowing the two domains to interact freely with RNA and liposomes.

Surprisingly, we observed that both the Ct and Nt are capable of interacting with liposomes and RNA in a Ca2^+^-dependent manner, and these interactions can occur simultaneously. This dual binding and its calcium-regulated hierarchy finely tunes ANXAll’s binding to RNA and lysosomes, promoting a large complex essential for overcoming transport steric hindrance. Moreover, our result showed that the p.D40G mutation, in the Nt domain, associated with ALS, displays destabilized interdomain interactions and bypass Ca2^+^ regulation, leading to aberrant aggregation. These insights advance our understanding of ANXAll’s role in neuronal RNA transport and its disruption in neurodegeneration, highlighting potential targets for therapeutic intervention.

## Introduction

Recent studies have shown that organelles are transported along microtubules by motor “vehicles” carrying “hitchhikers” like protein complexes or organelles(1,2). Tethers connect vehicles and hitchhikers, enabling rapid transport of materials such as mRNAs to dendrites or axon terminals. Annexin A11 (ANXA11) acts as a key tether, linking RNA granules to lysosomes via its N-terminal domain (Nt) binding RNA and its C-terminal domain (Ct) with lysosomes, facilitating passive transport of mRNAs within neurons(3).

ANXA11 is a 56 kDa calcium-binding protein belonging to the annexin family, which is present across all kingdoms except yeast (4). Humans have 12 annexins (A1–A11 and A13)(4). These proteins share a highly conserved Ct with four repeats, each capable of binding Ca^2^_+_ ions, composed of ∼70 amino acids forming five α-helices(5,6). The Ct adopts the form of a compact disk with a convex surface that interacts with cell membranes and a concave surface directed to the cytosol(7). In contrast, the Nt varies greatly in length and sequence among annexins, determining their specific functions(8); the Nt of ANXA1 is well-studied. It consists of 40 amino acids forming a helix–turn–helix motif (HtH) that interacts with the Ct. Calcium binding disrupts this interaction freeing the Nt to participate in protein-protein interactions(9). ANXA11, however, has a much longer Nt low-complexity domain (∼200 residues) rich in Gly, Tyr, and Pro, with a HtH helical motif around residues 38–59(10) predicted to have a comparable regulation mechanism to ANXA1(11,12)

ANXA11’s domain organization and tethering function may also contribute to neurodegenerative diseases like Amyotrophic lateral sclerosis (ALS), a condition affecting motor neurons. Recent studies identified mutations in ANXA11, such as p.G38R and p.D40G in the Nt, in both familial and sporadic ALS patients(12,13,14). These mutations are associated with abnormal aggregation of ANXA11 in motor neurons, which may disrupt cellular processes and promote neuronal degeneration. Although the exact mechanisms remain unclear (15)it is hypothesized that these mutations impair the tethering of RNA granules and lysosomes, essential for mRNA transport and cellular homeostasis, potentially leading to neuronal stress. Several key questions need to be addressed to deepen our understanding of this process and gather essential information to link the RNA transport to the ALS pathophysiology: what are the specific roles of the ANXA11 Nt and Ct in its tethering ability? Do these domains function independently or synergistically to mediate interactions with RNA granules and lysosomes? Are these two domains capable of directly interacting with each other, as established for other annexins such as ANXA1(9)? Would this interaction be calcium regulated? How these domains interplay will modulate the RNA and lysosome binding?

This study presents, for the first time, a detailed molecular mechanism underlying the interplay between the two ANXA11 domains, RNA, and liposomes. The results demonstrate that ANXA11 facilitates passive RNA transport by coordinating interactions with liposomes and RNA, with calcium serving as a crucial regulator. Additionally, our results reveal that the ALS-associated ANXA11 Nt-D40G (Nt_D40G_) mutation disrupts interdomain interactions between the Nt and Ct bypassing calcium regulation, leading to abnormal protein aggregation. These insights enhance our understanding of ALS pathology and could inform the development of therapies aimed at stabilizing ANXA11 interactions, restoring normal RNA transport, and mitigating neurodegeneration

## Results

### Structural characterization and calcium ions binding properties of ANXA11 Ct

Before testing ANXA11’s Nt and Ct functions, we confirmed their proper folding. Circular dichroism (CD) analysis of the Ct, without the thioredoxin carrier, revealed typical α-helix signals, with minima at 210 and 222 nm (Fig 1a, black line). K2D2 estimated about 70% α-helix content, aligning with other annexins(16,17,18). The Nt is mostly disordered with few structured elements(11).

**Fig. 1.**
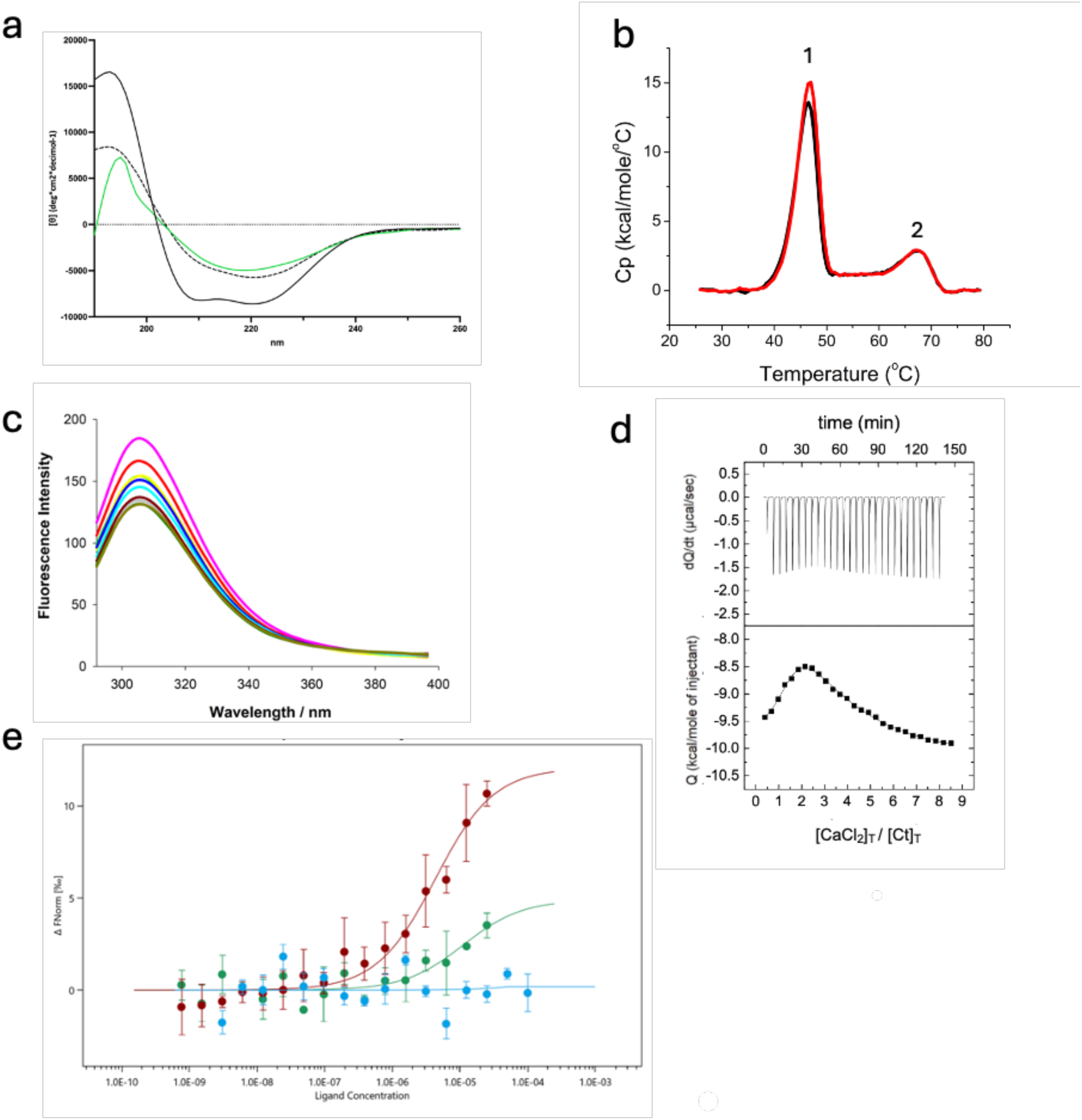
Calcium engagement with ANXA11 Nt and Ct governs its interdomain. **a** Far-UV spectra of Ct: black, 5°C; green, thermal unfolding at 75°C; dotted, cooling to 5°C. **b,** DSC thermogram of Ct in absence (black) and presence (red) of Ca^2+^; first derivative highlights transition midpoint (T_m_ = 46.53°C without Ca^2+^, 46.97°C with Ca^2t^). **c,** Fluorescence spectra of Ct (6 µM) with increasing CaCL_2_ (0-32 µM, λ_ex_ = 277 nm, 298 K, pH 7.45). **d,** ITC of Ca^2+^ (540 µM) binding to Ct (13.5 µM) at 25°C; top, thermogram; bottom, binding isotherm fitted with two-site model. **e,** MST quantification of C-N interaction: normalized fluorescence of labeled Ct titrated with unlabeled Nt at 20°C; fitting with MO Affinity Analysis v3.0.5.

We heated the Ct to 75°C, causing a broad minimum in the CD spectrum, indicating protein aggregation (Fig. 1a, green line). Cooling to 5°C did not restore the original spectrum (Fig. 1a, dotted line), suggesting irreversible aggregation. Moreover, Differential Scanning Calorimetry (DSC) for the combined protein (Fig 1b, black line) and thioredoxin alone (Fig. S1) as a control that showed a single peak at 70°C. The Ct showed two endothermic peaks: pick1 at 46.5°C for the Ct and pick2 at 68.5°C attributed to Thioredoxin. A similar test for the Nt showed no clear thermal transition, which was expected because the Nt is intrinsically disordered (data not shown)(11).

### ANXA11 Ct domain binds calcium

To investigate and quantify the Ct ability to bind Ca^2^_+_, we performed DSC experiments where the Ct thermal transition was measured in the presence and absence of Ca^2^_+_ in 10 fold molar excess with respect to ANXA11 Ct. The experiment in presence of Ca^2^_+_ showed that the melting temperature (Tm) of the Ct domain was slightly elevated (46.97°C; Fig. 1b, pick 1 red line) compared to the Tm in the absence of calcium (46.53°C; Fig. 1b, peak 1, black line). To assess calcium binding to the Ct domain, we performed DSC experiments measuring its melting temperature with and without Ca^2^_+_ (10-fold excess with respect to Ct). Ca^2^+ caused a slight increase in Tm from 46.53°C to 46.97°C (Fig. 2b, pick 1, red and black line respectively) and raised enthalpy from 107 to 113 kcal/mol, indicating a Ca^2^+ binding, likely through interactions with outer loops. This effect was specific to Ct, as calcium did not alter Thioredoxin’s thermal behavior (Fig. 1b, pick2, read and black line). To further analyze the calcium binding properties of Ct, we conducted fluorescence experiments with increasing Ca^2^_+_ (Fig. 1c). The addition of Ca^2^_+_ induced a quenching up to 25% of the fluorescence intensity at 306 nm supporting that Ca^2^_+_ directly binds to Ct.

**Fig. 2.**
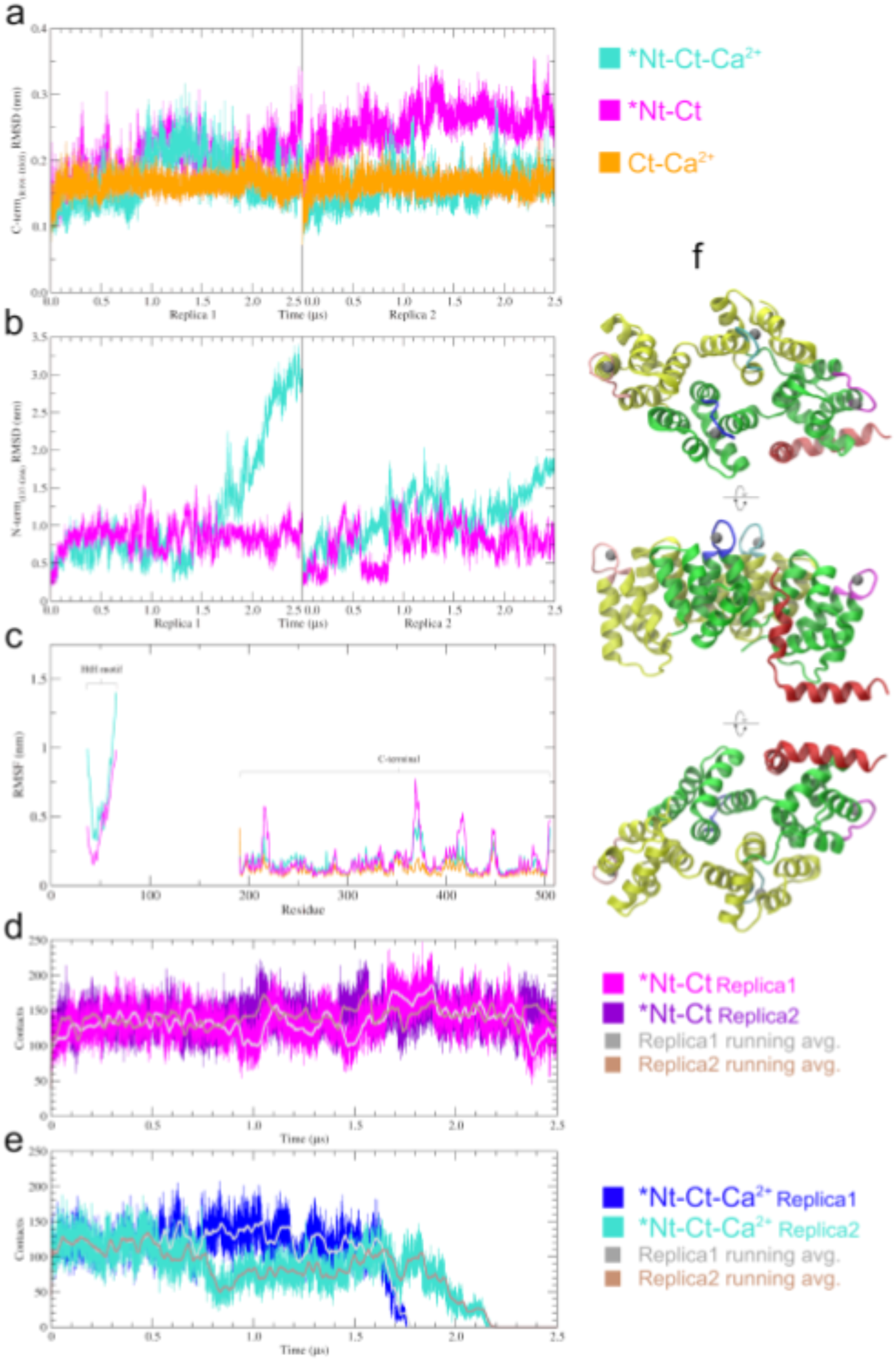
MD analysis of ANXA11 Cl and Nt HtH motif with and without Ca^2+^. **a.** RMSD analysis of ANXA11 C-terminal domain for the *Nt-Ct-Ca^2+^(cyan), the *Nt-Ct (magenta), and the control system. Ct-Ca^2+^ (orange). **b.** RMSD analysis of ANXA11 N-teπninal HtH motif when bound to Ct, with and w ithout Ca^2+^ (cyan and magenta, respectively). **c.** RMSF analysis of Nt HtH motif and Ct domain for the three systems: *Nt-Ct-Ca^2+^ (cyan), *Nt-Ct (magenta), and Ct-Ca^2+^ (orange, control); for the last one no HtH is present, d, C’t;Nt contact analysis performed for the *Nt-Ct system, in magenta and violet the contact count along Replica1 and 2, respectively, with the related running averages (grey and light-brown). **c,** Ct:Nt contact analysis performed for the *Nt-Ct system ’Nt-Ct-Ca^2+^ system, in blue and cyan the contact count along Replica 1 and 2, respectively, with the related ninning averages (grey and brown). **f**, ANXA11 Ct cartoon structure (top, bottom and side views, starting conformation of the *Nt-Ct-Ca^2+^ simulations) in the presence of both the Nt HtH motif and the four Ca^2+^ ions. The repeats forming the Ct domains are coloured green and yellow, the HtH motif in red. The AB loops are highlighted in pink, blue, cyan and magenta. The Ca^2+^ are show^r^n as grey spheres.

To quantify Ct interaction with Ca^2^_+_, we used Isothermal titration calorimetry (ITC) that provided accurate measurements of binding affinity, enthalpy, and stoichiometry. The titration with Ca^2^+ revealed biphasic thermograms (Fig. 1d), indicating two distinct binding processes and distinct binding sites. Four binding sites were identified: two high-affinity with dissociation constant (K_d_) = 1.8 ± 0.4 µM and two moderate-affinity with K_d_ = 38 ± 8 µM. These two calcium-binding sites have different strengths and thermodynamic behaviors. Indeed all sites bind calcium mainly due to entropy (-TΔSı = -8.3 kcal/mol and -TΔS2 = -14.9 kcal/mol), with little or no benefit from enthalpy (ΔHı = 0.5 kcal/mol and ΔH2 = 8.9 kcal/mol). The discovery of two K_d_ for Ca^2^_+_ binding to Ct suggests that the four sites can be divided into two groups. Although annexins’ ability to bind calcium is well known(19), to our knowledge, this study provides the first direct quantification of Ca^2^+ affinity for ANXA11 Ct.

### Nt and Ct domain interaction is calcium dependent

After characterizing the structure and calcium-binding ability of Ct, we wanted to investigate the possible intra-molecular interaction between Nt and Ct of ANXA11. Such intramolecular interactions have been already described for ANXA1(11,20), but never for ANXA11 with long Nt.

To test this interaction, we used microscale thermophoresis (MST). We labeled ANXA11 Ct with a fluorescent dye and gradually added increasing amounts of the unlabeled Nt (0.75 nM - 25 µM), testing both with and without different Ca^2^+ levels. We observed changes in the fluorescence signal indicating that the Nt and Ct bind with a K_d_ of about 4.3 ± 2.4 µM without Ca^2^_+_ (Fig. 1e, red circles). Upon adding Ca^2^_+_ at 15 and 30 times with respect to Ct concentration the binding weakened significantly. At 15 times Ca^2^_+_ excess, the K_d_ increased to around 11.2 ± 4.6 µM, and at 30 times excess, the binding was no longer detectable (Fig. 1e, green and blue circles respectively). These findings indicate that Ca^2^_+_ disrupts the interaction between the two domains of ANXA11, suggesting that it negatively influences their intramolecular interaction. To provide a detailed atomistic and dynamic representation of the Nt:Ct interaction, both in the presence and absence of Ca^2^_+_ ions, we conducted two independent 2.5 *µs* long molecular dynamics (MD) simulations for the following models: *Nt-Ct-Ca^2+^ (the Ct domain, R191-D505, plus a peculiar helix-turn-helix (HtH) motif of the Nt, I37-G66, plus four bound-Ca^2+^ ions, grey spheres), *Nt-Ct (same model as previous one, without Ca^2+^ ions), Ct-Ca^2+^ (control model including the Ct domain and the four Ca^2+^ ions).

The root mean square deviation (RMSD) analysis reports on the stability of the Ct (Fig. 2a) and the Nt HtH motif (Fig. 2b) in different conditions. In particular, in the presence of Ca^2+^ ions (Fig. 2a, *Nt-Ct-Ca^2+^, cyan curve) the Ct is quite stable in absence or presence of the Nt HtH motif (Fig. 2a, Ct-Ca^2+^ and *Nt-Ct-Ca^2+^ systems, cyan and orange curve, respectively). On the contrary, in the *Nt-Ct system (Fig. 2a, magenta line) the RMSD variation reached values above 0.3 nm, highlighting a higher flexibility.

Indeed, in the absence of Ca^2^+ ions, Ct residues in the AB loops (K214-E220, K286-E292, E369-E376, R445-D451 in magenta, blue, pink and cyan respectively in Fig. 2f) fluctuate much more than in the other two systems, as shown by the root mean square fluctuation (RMSF) analysis reported in Fig. 2c. The behaviour of the Nt HtH motif is quite peculiar: it remains quite stable in the absence of Ca^2^_+_ but exhibits huge flexibility in its presence. Indeed, the RMSD of the HtH motif in *Nt-Ct system increases significantly in the second part of the simulations, with respect to that in the *Nt-Ct-Ca^2+^ system (Fig 2b, magenta and cyan curves, respectively). Accordingly, also the RMSF shows higher values (Fig 2c) The visual inspection of the MD trajectories show a complete unbinding of the Nt HtH fragment from the Ct, thus suggesting that the presence of Ca^2+^ unfavour Nt:Ct interaction (see Fig. S4a, Videos 1A/1B). On the contrary, in both replicas of the *Nt-Ct simulations the final structure revealed a bound conformation, with the HtH motif still interacting with the Ct domain (Fig. S4b, Videos 2A/2B). This is supported by the counting of the Nt:Ct contacts reported in Fig. 2d/e. Indeed, while in the *Nt-Ct system (Fig. 2d), the number of contacts remain stable during the simulation, in the *Nt-Ct-Ca^2+^ one (Fig. 2e) we observe a large drop of interactions eventually reaching zero when the two domains are completely separate. The control simulations of the Ct-Ca^2+^ confirmed the persisting ions-binding at the AB loop sites (Figs. S2j and S4c).

Our results strongly suggest that ANXA11 exists in two conformations: a compact “close” form where the Nt and Ct interact intramolecularly, and an “open” form where this binding is lost, likely allowing for interaction with other molecules. Importantly, this conformational change is Ca^2+^-dependent and appears to be a key feature of ANXA11’s function, facilitating interactions with RNA and lysosomal membranes. This represents the first experimental confirmation of such an intramolecular interaction within the annexin with long Nt.

### ANXA11 Nt and Ct domain binds lipid-vesicles in a calcium dependent manner

ANXA11’s Nt and Ct intramolecular interaction, its calcium regulation, and recent fmdings(3) on its hierarchical binding to RNA granules and lysosomes offer valuable new insights into how ANXA11 functions in both normal and disease states. Building on the idea that the Ct is involved in lysosomal membrane binding(3), we investigated this interaction *in vitro* to understand its molecular basis. Additionally, to comprehensively explore the functions of ANXA11 domains, we examined whether the Nt could also bind to lipid membranes, similar to Ct.

To achieve this, we conducted pull-down assays using purified Ct and Nt incubated with liposomes (LipoA)(21). These synthetic lipid vesicles serve as an excellent *in vitro* model for studying protein-membrane interactions(21). To investigate Ca^2^_+_ dependence, we tested increasing calcium concentrations.

Our results demonstrate, for the first time, that the Ct domain binds LipoA in a Ca^2+^-dependent manner, with binding efficiency increasing proportionally to calcium concentration (Fig. 3a, lanes 3-6). The proportion of Ct bound to LipoA ranged from 78.9% to 88.8% at lower calcium concentrations (l:5 and l:l0 Ct/Ca^2+^ ratio) and increased to 96.3% and l00% at higher calcium concentrations (l:l5 and l:20 Ct/Ca^2+^ ratio). These percentages were compared to a baseline of l00% at the highest calcium concentration (l:20 Ct/Ca^2+^ ratio).

**Fig. 3.**
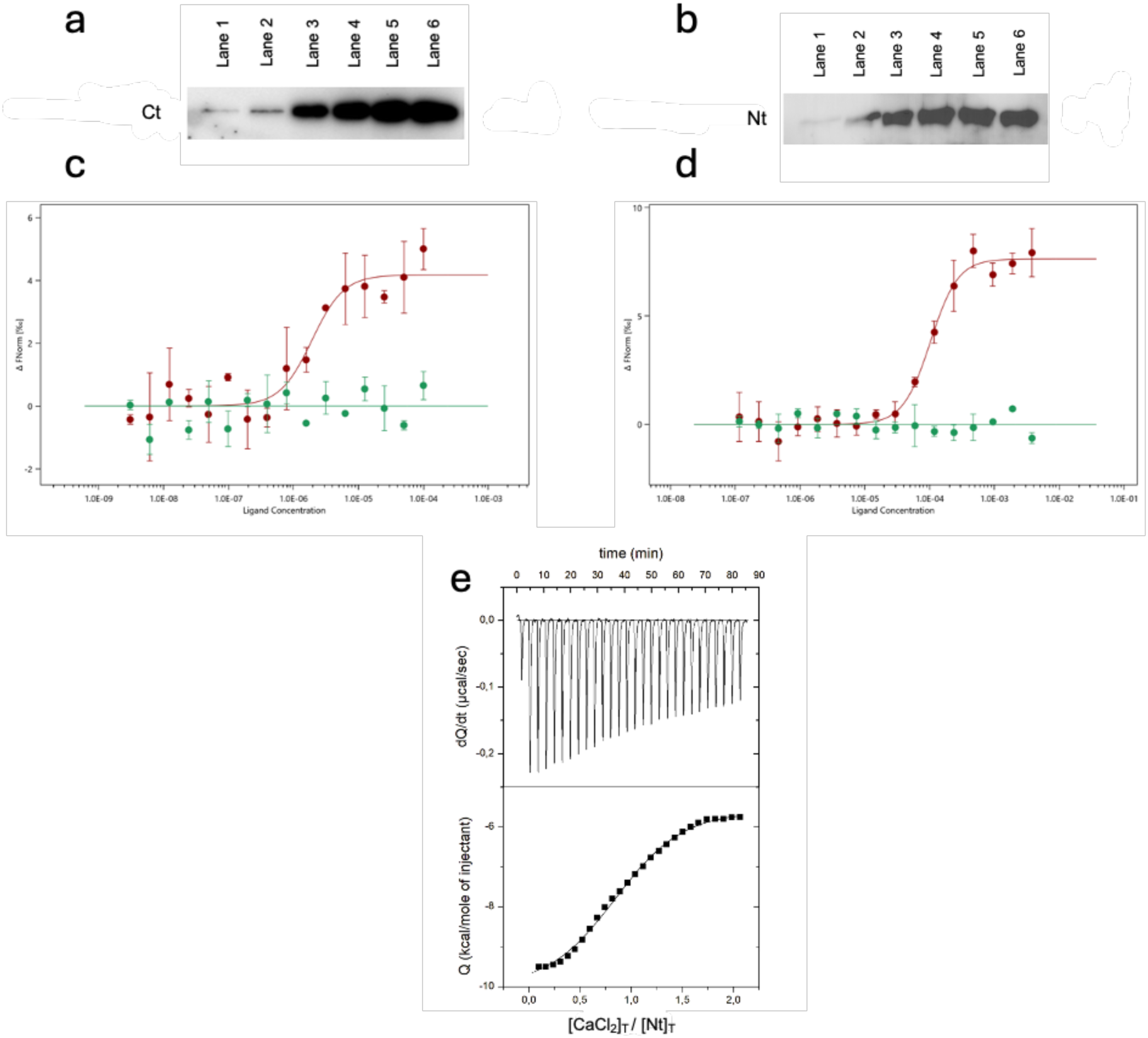
Calcium-regulated interactions of isolated ANXA11 domains with lipid membranes. **a**, Western blot pulldown assay showing binding of Ct to LipoA under a calcium gradient. Lane 1: 7.5 mM EDTA; Lane 2: no calcium; Lanes 3-6: increasing molar equivalents of Ca^2+^ (5, 10, 15, 20 ME). **b**, Western blot pulldown assay showing binding of Nt to LipoA under a calcium gradient. Lane 1: 7.5 mM EDTA; Lane 2: no calcium; Lanes 3­6: increasing molar equivalents of Ca^2+^ (5, 10, 15, 20 ME). **c**, MST quantification of labeled Ct titrated with 150 run LipoB in the presence of a lO× molar excess of calcium (red) or 7.5 mM EDTA control (green). Measurements at 20 °C, 10% excitation, 40% MST power; cold and hot regions from -1.0 to 0 s and 1.58-1.36 s. **d**, MST quantification of labeled Nt titrated with 150 run LipoB in the presence of a lO× molar excess of calcium (black) or 7.5 mM EDTA control (grey). Measurements at 25 °C, 100% excitation, 40% MST power; cold and hot regions from -1.0 to 0 s and 4-5 s. **e**, Surface charge density of the Ct without calcium: AB loops partially hidden, negatively charged residues at calcium-binding sites less exposed, while opposite face shows a generally negative patch. **e**, ITC of Ca^2+^ (100 µM) binding to ANXA11 Nt (10 µM) at 25 °C; top, thermogram; bottom, binding isotherm fitted with a one-site model.

This variation can be attributed to the differing Ca^2+^ affinities of the four calcium-binding sites, as identified by the ITC experiment. Higher Ca^2+^ concentrations facilitate full occupancy of the Ct calcium-binding sites, enabling this domain to function fully. Negligible binding was observed when the Ct was tested alone or with EDTA (Fig. 3a, lanes 2 and 1 respectively). This shows that calcium is essential for the Ct domain to bind to lipids.

Strikingly, our results reveal for the first time that the Nt can bind LipoA in the presence of Ca^2+^ (Fig. 3b, lanes 3-6), without showing the concentration dependence seen with the Ct. No binding occurred with EDTA (Fig. 3b, lane 1). Interestingly, the Nt weakly binds LipoA without calcium (Fig. 3b, lane 2). This novel finding unexpectedly suggests that the Nt domain can interact with LipoA, a possibility not previously hypothesized and it is particularly surprising that Ca^2+^ is necessary for this interaction.

### Quantification of the Calcium-dependent interaction between ANXA11 Nt and Ct and lipid vesicles

To investigate direct binding between the Ct and Nt with LipoA, we used 150 nm extruded unilamellar vesicles (LipoB). This approach ensured more uniform vesicle size and minimized aggregation, allowing for accurate quantification of binding. Specifically, we carried out MST measurements in the presence of 10 and 5 nM of Ct and Nt respectively labelled with RED-NHS fluorescent dye. Increasing concentration of the label-free LipoB was added in both experiments in presence of 10 molar excess of calcium with respect to the two domains. The fluorescence signal changes showed that the Ct interacts with the lipid vesicles (LipoB) producing an EC_50_ of 1.97 ± 0.88 μM (Fig.3c, red circles), in agreement with what is reported in literature(22). On the other hand, the fluorescence changes during the experiment where Nt interacts with LipoB showed an EC_50_ of 100 ± 20 µM (Fig. 3d, red circles). As a control we tested the interaction for Ct and Nt in the presence of EDTA, which completely prevented binding (Fig. 3c/d, green circles respectively). These results support the earlier pull-down experiments, reinforcing the hypothesis that the Ct and Nt require Ca^2+^ to promote their interactions with lipid vesicles.

Given the significant and unexpected influence of Ca^2+^ on ANXA11 Nt interactions with RNA and membranes, we investigated whether this effect was due to direct binding. ITC experiments showed a single high-affinity site per Nt molecule, with a 1:1 stoichiometry and a K_d_ of 2.4 ± 1 µM (Fig. 3e), confirming that the Nt domain directly binds Ca^2+^.

Ca^2+^ binding to the Nt is mainly entropy-driven, with minimal enthalpic contribution (ΔH = – 5.93 kcal/mol) and a favorable entropy term (–TΔS = –4.85 kcal/mol). This indicates Ca^2+^ may stabilize flexible regions and facilitate the domain’s interactions with RNA and membranes. These results offer a quantitative understanding of how calcium influences the Nt’s function and its role in regulating ANXA11 molecular interactions.

### Surface charge density analysis reveals Ca^2+^-induced conformational changes

To further investigate the Ca^2+^ effects on the Ct structure we performed the surface charge density analysis. We have reported in Fig. 4a and 4c the charge density maps of the *Nt-Ct and Ct-Ca^2+^ simulations most populated cluster conformation (i.e. the centroid that best represents the highly populated, similar MD-sampled conformations; see SI for details). By comparing the Ct concave-side opposite to the one binding the Ca^2+^ ions in the two systems, we observed, in the presence of Ca^2+^, a shift towards a neutral/partially positive exposed surface (Fig. 4b/right), at the expense of the negatively charged residues that were exposed in the absence of Ca^2+^ (Fig. 4a/right). The Ca^2+^ driven conformational change led to the hiding of the previously exposed negative charges, thus the Ca^2+^ might promote the exposure of a suitable surface (on the opposite side with respect to the same Ca^2+^ binding sites) that seems more prompt to interact with other partners. Further, the mapping epitope analysis was in agreement with this finding: concerning the Ct-Ca^2+^ simulations in the presence of Ca^2+^, it actually reported the appearance of a region that is likely prone to make an interaction (in red, pointed by the arrow in Figure 4d), possibly with negatively charged interactors; the same region was instead idle in the *Nt-Ct representative conformation (Figure 4c) in the absence of Ca^2+^ ions (further details in SI and Fig. S5). According to Alphafold 3(23) predictions (Fig. S3) the Ct Ca^2+^-binding side (exposing the AB and CD loops; convex side) could be involved in contacting RNA, thus leading us to hypothesize that the opposite side of the Ct (concave side) could likely be responsible for interacting with liposomes.

**Fig. 4.**
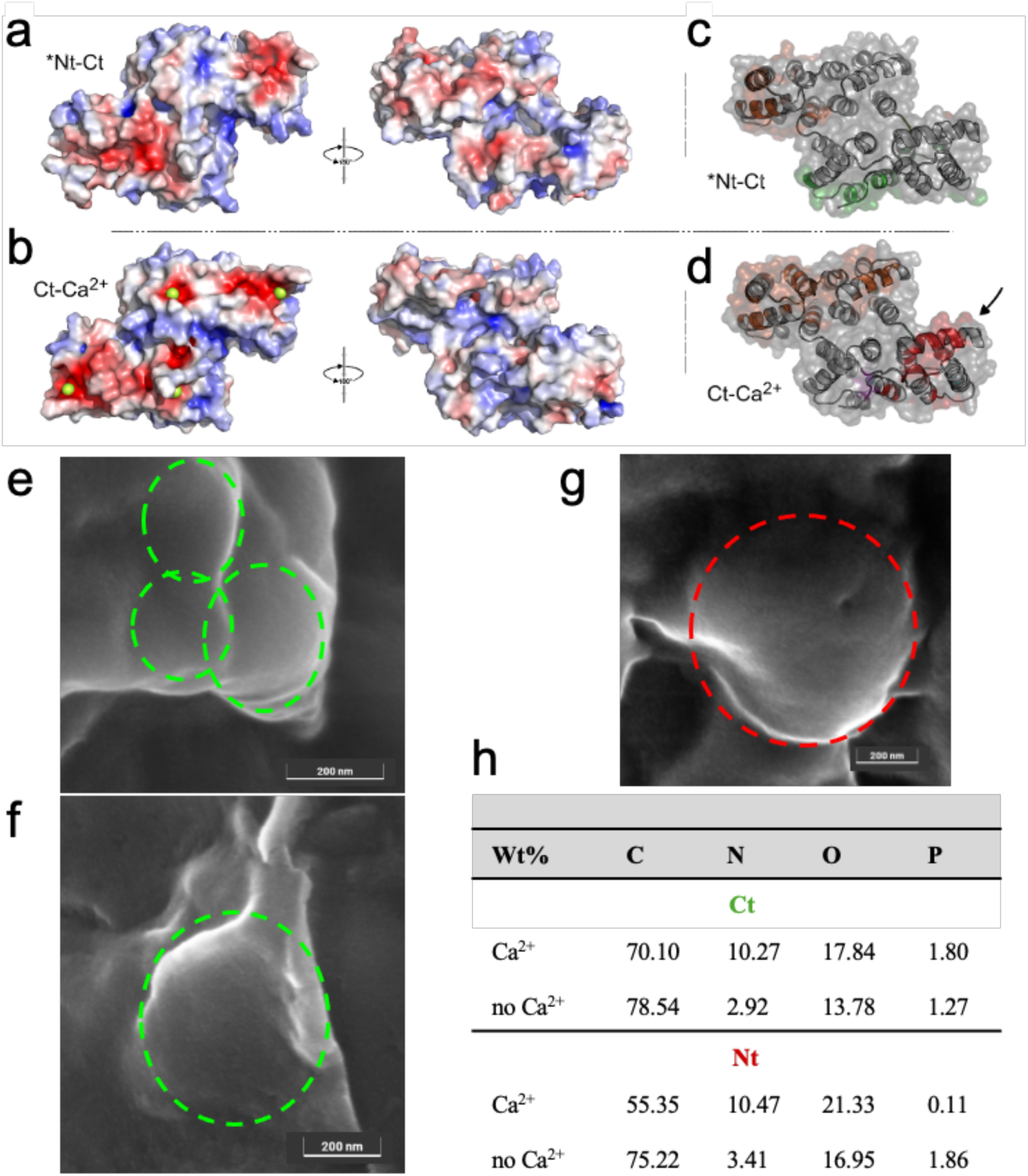
Molecular dynamics, ANXA11 Ct surface and epitope analysis with calcium, and FE-SEM/EDS of ANXA11 isolated domains with lipid vesicles. **a**, Surface charge density of the Ct without calcium: AB loops partially hidden, negatively charged residues at calcium-binding sites less exposed, while opposite face shows a generally negative patch. **b,** Surface charge density of the Ct in the presence of four Ca^2+^ ions (green spheres): calcium-binding loops well structured, providing stable cation interaction, while the opposite side shows neutral or positive regions suitable for interaction with negatively charged lipids. **c**, Epitope mapping of the centroid structure of the Nt-Ct most populated cluster without calcium. The surface opposite to the calcium-binding site is shown in the foreground. **d**, Epitope mapping of the Ct-Ca^2+^ system; a new epitope (red, pointed by the arrow) appears in the presence of calcium, visible on the surface opposite the calcium-binding site. **e**, Representative FE-SEM image of ANXA11 Ct (20 µM) + lipid vesicles 200 µM in the presence of a CaCL_2_ solution (400 µM). **f**, Ct (20 µM) + lipid vesicles 200 µM in the absence of a CaCL_2_ solution (400 µM). **g**, Nt (20 µM) + CaCL_2_ solution (400 µM) + lipid vesicles (200 µM) + CaCL_2_ solution (400 µM). **h**, Wt% of the different elements monitored by EDS in the presence or absence of Ca^2+^ for the Ct (green) and Nt (red) fragments. Dashed lines show the morphology of the LipoA vesicles. Instrumental magnification: 345000×, 245000× and 303000×, respectively.

### Visualization of protein-lipid vesicle interactions and calcium effects using electron microscopy

Finally we decided to perform FE-SEM experiments with Ct and Nt in the presence and in the absence of Ca^2+^. This will allow detailed visualization of surface morphology, structure of the lipid vesicles (appearing as globular particles ranging between 100 and 250 nm) and check if protein layers are coating them. Figure 4e presents a FE-SEM image of the Ct with lipid vesicles in the presence of Ca^2+^. These vesicles are covered by a layer likely formed by the ANXA11 Ct. This layer is observed in Figures 4e and 4g and its presence is further confirmed by EDS analysis shown in Figure 4h. A similar layer and morphology of the lipid vesicles were seen with the Nt (Fig. 4f).

In more details, added Ca^2+^ induced the formation of vesicles clusters covered by the Ct. This suggests that Ca^2+^, interacting with the Ct, encourages vesicles to clump together (Fig. 4e). As for this sample, these vesicle clusters were hardly observed without Ca^2+^ (Fig. 4f). Conversely, the Nt did not show this effect, as the shape of the vesicles with and without Nt was the same (Fig. 4g).

EDS spectra were collected from different areas of the Ct and Nt samples. This helped us examine the surface elemental composition and determine the location of the Ct and Nt on the lipid vesicles at the nanometer scale. The EDS results show that the main elements in the samples are carbon (C), nitrogen (N), oxygen (O), and phosphorus (P), which matches the expected composition of the Ct (10.27 wt% vs. 2.92 wt%) and Nt (10.47 wt% vs. 3.41 wt%) (Fig. 4h). Moreover, figure 4h shows that the elemental composition of both Ct and Nt samples changes with Ca^2+^ concentration. Specifically, the amount of nitrogen (N) detected by EDS is higher in the presence of Ca^2+^, suggesting that the outer surface of the lipid vesicles is enriched in N. This supports the idea that Ca^2+^ interacts with both Ct and Nt, strengthening their bond with the lipid particles. The changes in composition were more pronounced in the Ct (around 75% of the examined regions) compared to the Nt (about 5%) further respecting the differences with their relative affinity (i.e. K_d_) with Ca^2+^.

### Nt and Ct domains bind to RNA in calcium dependent manner

Although *in vivo* studies have confirmed that the ANXA11 Ct interacts exclusively with lysosomes and the Nt with RNA(3), mounting evidence suggests that the Ct can also bind specific RNA species(11,22). Additionally, it has previously been demonstrated that the ANXA11 Nt is essential for binding to RNA granules in neurons(3). To explore whether the Nt and Ct can directly bind RNA and to evaluate the effect of Ca^2^_+_ on these interactions, we conducted a series of *in vitro* pull-down assays.

These experiments utilized purified recombinant Nt and Ct domains and employed poly-U agarose beads (poly-U) as a synthetic RNA mimic. The assays were conducted under conditions both with and without the presence of Ca^2+^ to assess its impact on binding affinity and specificity. Nt and Ct were incubated with poly-U at various Ca^2+^ concentrations (1:5, 1:10, 1:15, and 1:20, relative to the protein concentration). Under these conditions, we observed that the Nt and Ct are effectively bound to the poly-U across all Ca^2+^ tested ratios (Fig. 5a and 5b, lane 3 and 6 respectively). Interestingly, when we performed the binding test without adding Ca^2+^, Nt and Ct were still slightly bound to poly-U (Fig. 5a/b, lane 2). This suggests that leftover Ca^2+^, possibly from purification, might be enough to support some binding, similar to what we saw with liposomes. As a negative control, we used EDTA and found negligible or no binding for both domains (Fig. 5a and 5b, lane 1).

**Fig. 5.**
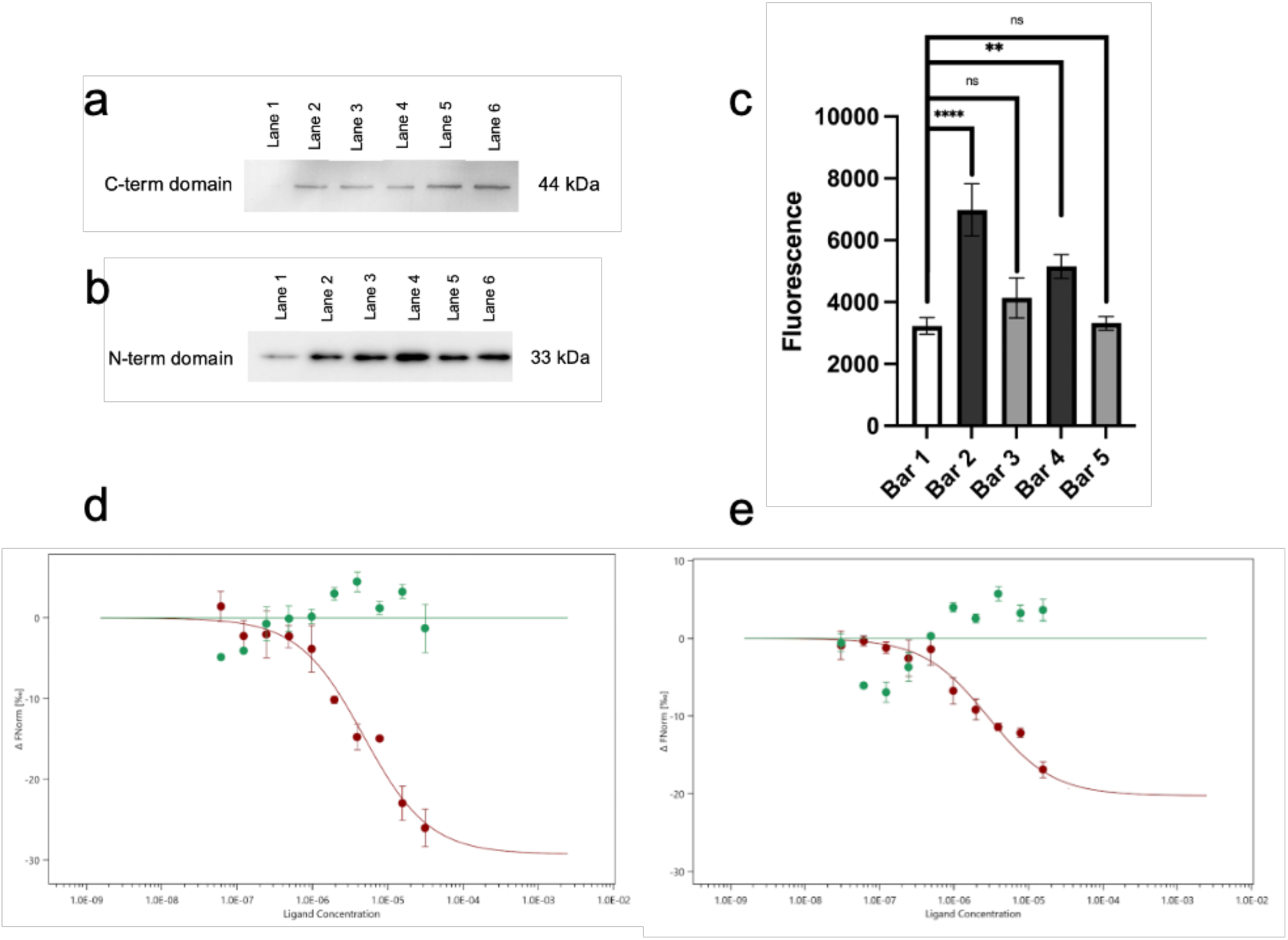
Calcium-regulated RNA interactions of isolated ANXA11 Nt and Ct domains. **a**, Western blot pulldown assay: ANXA11 C-term binding to poly-U agarose resin under a Ca^2+^ gradient (Lane 1 EDTA, Lane 2 no Ca^2+^, Lanes 3-6 with 5-20 ME Ca^2t^). **b**, Western blot pulldown assay: ANXA11 N-term binding to poly-U agarose resin under a Ca^2+^ gradient (Lane 1 EDTA, Lane 2 no Ca^2+^, Lanes 3-6 with 5-20 ME Ca^2+^). **c**, Pulldown fluorescence assay: RNA 6-FAM retained on NiNTA resin with Ct in presence of Ca^2+^ (Bar 2, **** p < 0.0001, unpaired two-tailed t-test) or EDTA (Bar 3, ns), RNA 6-FAM retained on NiNTA resin with Nt in presence of Ca^2+^ (Bar 4, ** p < 0.01) or EDTA (Bar 5, ns). **d**, MST quantification: Ct binding to 10-mcr poly-U RNA with Ca^2+^ (red) or EDTA (green). **e**, MST quantification: Nt binding to 10-mer poly-U RNA with Ca^2+^ (red) or EDTA (green).

Remarkably, our data provide the first direct evidence that the binding of the Nt and Ct to RNA is calcium-dependent. The presence of calcium markedly enhances this interaction *in vitro,* unveiling a previously unrecognized regulatory mechanism that may influence ANXA11’s function in cellular contexts.

Additionally, a reverse pull-down assay confirmed Nt and Ct binding to RNA. We immobilized these domains on Ni-NTA resin (NiNTA) and incubated with 6-FAM-labeled poly-U 10-mer. Fluorescence changes were used to measure the interaction. Figure 4c summarizes the results. Labeled RNA alone added to NiNTA gave a fluorescence intensity of 3229 ± 263 (Fig. 5c, bar 1). With Ca^2^_+_, signals increased to 5152 ±389 when Nt was present (Fig. 5c, bar 4) and 6979 ± 845 when Ct was added (Fig 5c, bar 2). Adding EDTA reduced signals to near control levels (3317 ±219 and 4,139 ±650 respectively Fig. 5c, bar 3 and 5). These signals were within the control standard deviation. These results indicate that Ca^2+^ is crucial for RNA binding by both domains supporting our poly-U agarose pull-down findings.

### Quantification of the Ca^2^_+_ dependent interaction between ANXA11 Nt and Ct with RNA

We further quantified the interaction between the Ct and Nt with poly-U 10-mer using MST where 5 nM of labeled Ct and Nt (Kit RED-tris-NTA) were used in the presence of increasing concentrations of unlabeled RNA, ranging from 7.63 nM to 31,25 μM. All measurements were performed in the presence of a 10-fold molar excess of calcium relative to the domains . The changes in fluorescence signal provided evidence of Ct-RNA interaction (Fig. 5d, red circles) and Nt-RNA interaction (Fig. 5e, red circle), resulting in an estimated K_d_ of 3.55 ± 0.30 µM and 4.18 ± 0.80 µM respectively. To confirm Calcium’s role in Ct and Nt interactions with RNA, the experiments were repeated with EDTA. In this condition the RNA interaction with Nt and Ct is abolished (Fig. 5d and 5e, green circles), supporting that Ca^2+^ is essential for their interaction.

These results show that Ca^2^_+_ is necessary for the Ct to bind RNA, but the binding is not Ca^2^_+_ concentration dependent. More importantly, Ca^2^_+_ is clearly necessary for the Nt to bind RNA. This indicates a new way that calcium regulates ANXA11-RNA interactions, which could affect its function in cells

### ANXA11 Nt and Ct domains bind simultaneously to RNA and lipid vesicles in a calcium dependent manner

After independently analyzing the binding of the Nt and Ct to lipid vesicles and RNA, we tested if Nt and Ct can bind simultaneously to both ligands. Using a pull-down assay, we adsorbed either Nt or Ct to LipoA vesicles with excess Ca^2^_+_ (20-fold molar excess with respect to the domain), then added a fluorescent 10-mer RNA. Fluorescence measurements showed RNA binding.

Figure 6a (bars 2 and 3) shows that Ct and Nt bind RNA-6FAM in the presence of LipoA and Ca^2^_+_, as resulting by the significantly higher fluorescence observed (6961 ± 1139 and 6363 ± 815 arbitrary units, respectively) compared to a control (Fig. 6a, bar 1) where Nt and Ct were added only with LipoA and RNA-6FAM (1676 ± 323 arbitrary units). Statistical analysis confirmed that the fluorescence values of the Ct and Nt in the first reported conditions were significantly higher than the control (p = 0.0001), supporting the conclusion that both domains are capable of simultaneously interacting with lipid membranes and RNA in a calcium-dependent manner.

**Fig. 6.**
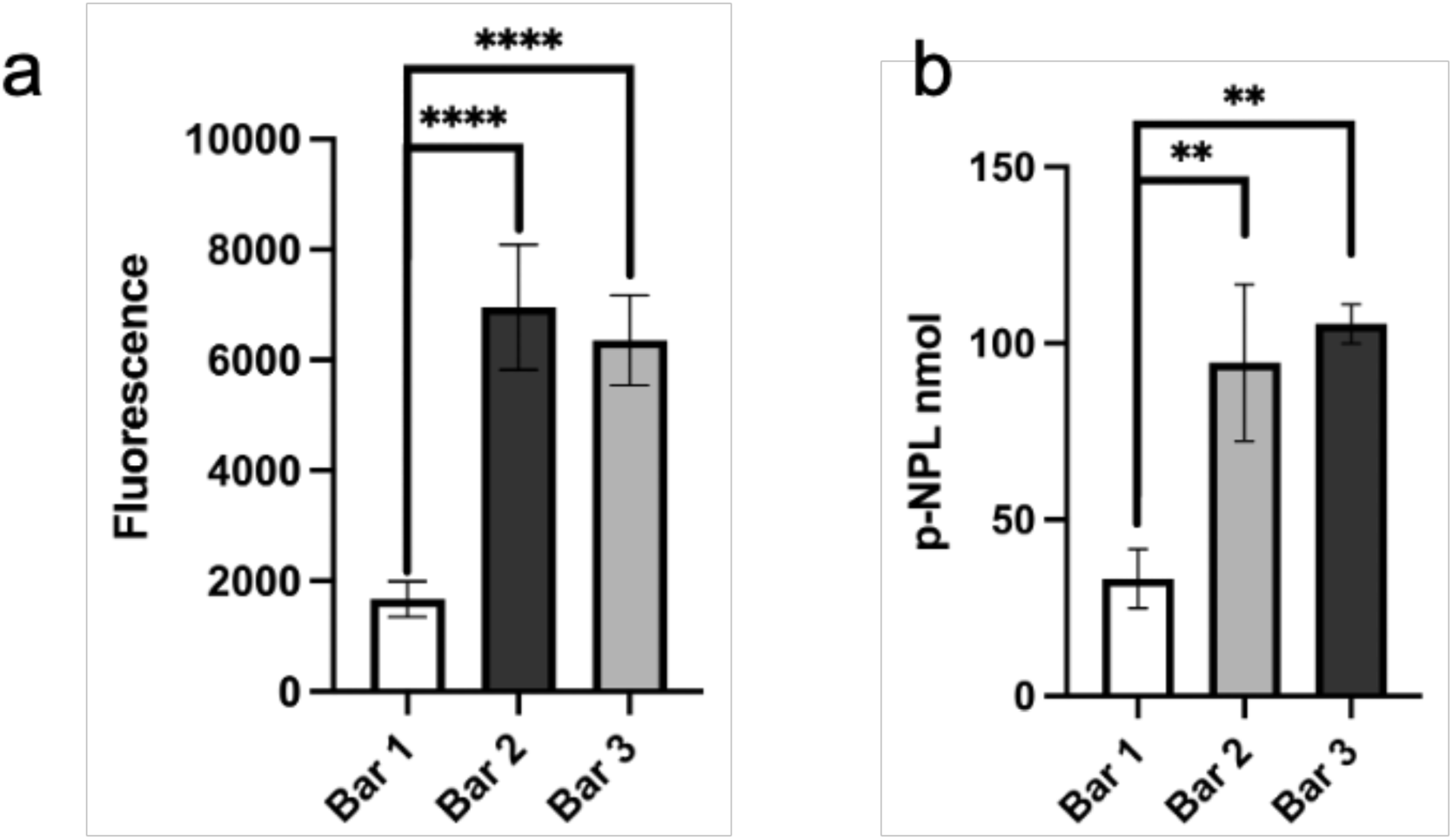
Simultaneous lipid and RNA engagement by ANXA11 Nt and Ct domains. **a**, Pulldown assay: fluorescence of ANXA11 domains and poly-U labeled with 6-FAM. Bar 1, LipoA + RNA 6FAM, Bar 2, LipoA + poly-U + C-tcrm + 20 ME of calcium; Bar 3, LipoA + poly-U 6FAM + Nt + 20 ME of calcium. (****) indicate a p-valuc < 0.0001 (unpaired two-tailed t-test), representing statistically significant differences between conditions. **b**, Interaction of micelles and Poly-U wth the ANXA11 Ct and Nt domains. Absorbance at 410 nm was measured to assess micelles binding via p-nitrophcnol release. Bar 1, poly-U + p-NPL (33.3 ± 8.33 nmol); Bar 2, poly-U + Nt + p-NPL + Ca^2^* 20 molar excess (94.50 ± 22.2 nmol); Bar 3, poly-U + Ct + p-NPL + Ca^2^’ 20 molar excess (105.50± 5.55 nmol). Measurements were performed in triplicate. (**) indicate a p-valuc < 0.01 (unpaired two-tailed t-test), representing statistically significant differences between conditions.

To confirm the combined interaction of Nt and Ct with RNA and lipid membranes, we bound Nt or Ct to poly-U. Subsequently, para-nitrophenol (p-NPL) micelles were added to facilitate the formation of a ternary complex. Figure 6b shows that p-NLP production significantly increased (∼3-fold) only when poly-U, p-NLP and Nt (Fig. 6b, bar 2) or Ct (Fig. 6b, bar 3) are present, while controls with only RNA or p-NPL showed no significant increase (Fig. 3b, bar 1). Additionally, a control experiment (protein domains + p-NPL, no poly-U) showed no precipitation artifacts (data not shown). This confirms that both domains can simultaneously associate with RNA and lipid membranes, supporting the formation of a stable complex.

### ANXA11 Nt_D4øG_ mutant disrupts the binding with Ct

To understand the role played by ANXA11 in ALS and gather information on the role played by ANXA11 interdomain interaction and within RNA transport, we decided to study the mutation Nt_D40G_ (carrying a single Asp-to-Gly substitution at position 40) identified in ANXA11’s Nt domain and found in ALS patients. This mutation is located right at the binding surface between the Nt and Ct domains of ANXA11, allowing us to characterize the binding properties of the Nt p.D40G mutant with the Ct, both with and without Ca^2^_+_. These findings are important for understanding how this disease-associated mutation affects ANXA11 physiological function.

We performed MST measurements using 10 nM labeled Ct, titrated with increasing (0.75 nM– 25 µM) unlabeled Nt_D4øG_. Fluorescence changes showed little to no binding (K >>20 µM, Fig. 7a, green). The wild-type Nt binding curve (K_d_ = 4.3 ± 2.4 µM) here is also shown (Fig. 7a, red) for comparison. To better understand how the Nt_D40G_ mutation within the HtH motif affects the Nt-Ct intradomain interaction, two MD replicas of *Nt_D40G_-Ct simulations were performed where the Ct did not present the four Ca^2^_+_ bound. This *Nt_D4øG_-Ct simulation was compared with the *Nt-Ct one carried out previously. This helped us assess the combined impact of the D40G mutation and calcium removal. As expected, RMSD and RMSF analyses of *Nt_D4øG_-Ct showed similar destabilization of the Ct domain’s loops observed in the *Nt-Ct, with deviations and fluctuations comparable to previous wild-type simulations without Ca^2^_+_ (Fig. 7b and 7d vs. Figs. 2a and 2c, magenta curves).

**Fig. 7.**
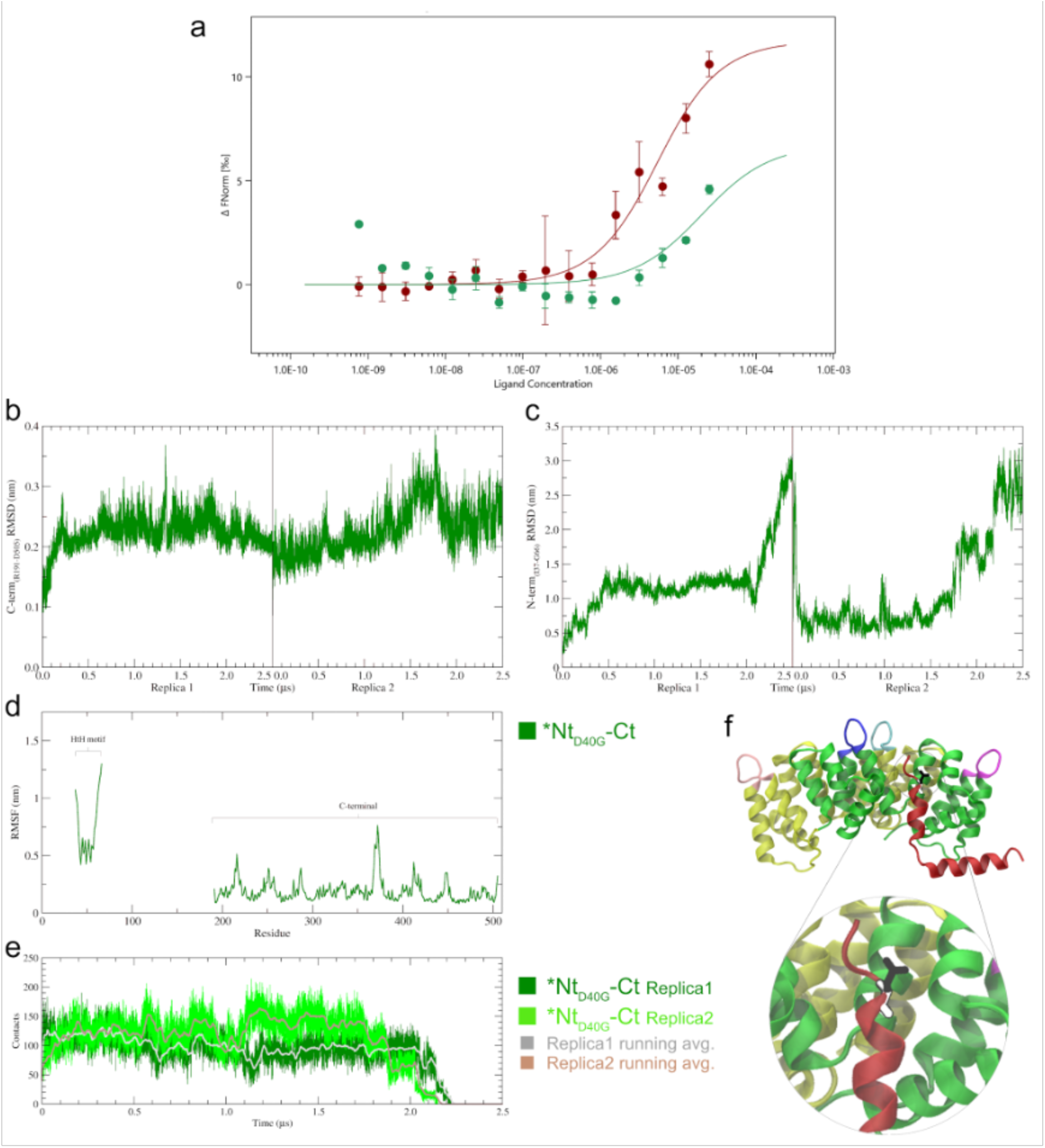
ANXA11 Ct-Nt interaction in the p.D40G mutant and analysis of the related simulations (*Nt_D40G_-Ct). **a**, MST quantification of Ct-Nt_D40G_ interaction: normalized fluorescence of labeled proteins titated with unlabeled Nt_D40G_; at 20°C (green), binding curve of the *wt* Nt shown for comparison (red); fitting with MO Affinity Analysis v3.0.5. **b,** Concerning the *Nt_D40G_-Ct simulations, RMSD analysis of the Ct domain (R191-D505) with respect to the starting structure; the Ct domain sampled distant conformation from the starting one due to the absence of Ca^2+^, like in *Nt-Ct simulations (Fig. 2a, magcnta). **c**, RMSD of the Nt HtH motif (I37-G66) carrying the p.D40G mutation, in the occurrence of an unbinding event in both replicas, HtH motif detaching from the Ct domain, a rapid increase of the deviation is observable. **d**, RMSF analysis of the *Nt_D40G_-Ct simulations: HtH motif and Ct domain arc pointed out; even in the absence of Ca^2+^ ions, due to the p.D40G mutation, the fluctuations of the HtH residues are large, in particular the L39-Q48 ones; large fluctuations in the Ct domain can be found at the four AB loop regions, and at residues D249-E257 (D helix) and Y279 that were initially pairing with the HtH. **e**, HtH/Ct contact analysis in the *Nt_D40G_-Ct simulations, in dark-and light-green the contact count along Replica 1 and 2, respectively, with the related running averages (grey and light-brown), both sampling the disruption of the interaction between the two domains at ∼2.2 µs. **f**, ANXA11 Ct structure (side view, starting conformation of the *Nt_D40G_-Ct simulations) with the interacting HtH (red) carrying the Asp-to-Gly mutation at position 40 (black/white licorice and zoomed in below); such an interaction will be lost during the simulations (Videos 3A and 3B).

Two new features appeared in the RMSF analysis of the D40G mutant. First, the HtH motif’s instability was similar to the *Nt-Ct-Ca^2^_+_ system (Fig. 7d vs. Fig. 2c, cyan). Second, fluctuations in the first helix (L39-Q48) of the HtH motif were even more pronounced than in other systems (Fig. 7d vs. Fig. 2c, cyan and magenta). Moreover, the residues D249–E257 and Y279 within the Ct showed increased flexibility not observed previously. These Ct residues normally interact with the HtH motif. The acquired flexibility of the mutated HtH, due to the missing Asp at position 40, appears to influence nearby residues on the opposite side of Ct (Fig. 7f structure).

The p.D40G mutation significantly destabilized the HtH-Ct interaction, even without Ca^2^_+_, unlike the *Nt-Ct simulations. By the end of *Nt_D40G_-Ct simulations, contacts dropped to zero (Fig. 7e), and RMSD rose to ∼3 nm (Fig. 7c) indicating the complete loss of intramolecular bonds. Final simulation-sampled conformations (Fig. S4d, Videos 3A/3B) confirmed the unbinding events. To our knowledge, this is the first experimental and *in silico* evidence demonstrating a role of this mutation within the ANXA11 function, highlighting that the p.D40G mutant bypasses the essential requirement of Ca^2^_+_ regulation based on the self­interaction between the Nt and Ct of ANXA11. This loss of regulation leads to the aberrant exposure of the Nt, even when it is not physiologically required, most likely resulting in protein precipitation.

## Discussion

To better understand the findings reported by Liao et al.(3) regarding the ability of ANXA11 to bind lipid membranes via Ct and RNA via Nt, we investigated these interactions and how they are influenced by Ca^2+^, by producing its Nt and Ct domains for individual analysis. These will reveal their individual binding affinities for RNA and lipid membrane and how they relate to the Nt:Ct interaction. Moreover, we produced the Nt_D40G_ mutant linked to ALS to evaluate how it could affect the Nt:Ct interaction.

Structural studies show the Ct is mostly helical, like other annexins, and its stability is increased by the Ca^2^_+_ binding probably through the interaction with surface loops. While the Ca^2^_+_-binding sites in many annexins are characterized(5,24), direct evidence of Ca^2^_+_ binding to annexins is scarce(19). We discovered by ITC that Ct has two sets of Ca^2+^-binding sites with high and moderate affinity, unlike ANXA4, which has uniform calcium binding across its four loops(19).

For the first time in annexins with a long Nt, we examined the role of Ca^2+^ in regulating interactions between the Nt and Ct, providing clear evidence of a direct interaction between these domains that is highly sensitive to Ca^2+^ levels. Specifically, increasing Ca^2+^ concentrations gradually weaken their interaction till the complete dissociation. This indicates that Ca^2+^ functions as a molecular switch: in the absence of Ca^2+^, the domains interact, thus remaining in a closed conformation; when Ca^2+^ is present, the interaction is disrupted, resulting in an open conformation where both domains are exposed to bind different partners.

MD simulations confirmed the experimental results adding detailed insights into the Ct:Nt interaction. Without Ca^2+^, the two domains bind stably. Ca^2+^ addition to the simulation disrupts the interaction both stabilizing the Ct structure and increasing HtH motif fluctuation. These opposite effects lead to the decoupling of their dynamics leading to their rapid dissociation. This ability to exist in two conformations is crucial for modulating cellular functions and enabling rapid responses to stimuli. Similar mechanisms have been observed in ANXA1, where calcium triggers the release of the short Nt to carry out its specific roles(9). This is the first documented Ca^2+^ regulatory mechanism for an annexin with a long Nt. It is tempting to suggest it may be a common feature across the annexin family.

Discovering Ca^2^_+_ ’s role in ANXA11 regulation led us to explore its link to ALS, focusing on the Nt_D4øG_ mutant linked to toxic protein aggregation(12). Our data showed that this mutation impairs Nt:Ct interaction when Ca^2^_+_ is absent, bypassing Ca^2^_+_ regulation. This persistent mutated-ANXA11 competence in keeping the open conformation likely promotes toxic aggregates linked to ALS neurodegeneration(12). Building on Ca^2^_+_’s role in controlling ANXA11’s switch between open and closed states, we explored how this regulation affects its interactions with membranes and RNA. Remarkably, for the first time, we demonstrated that Nt can directly bind Ca^2+^ and that this binding facilitates its Ca^2+^-dependent lipid binding. This unexpected discovery broadens the traditional understanding of annexin’s membrane association mechanisms. Moreover, our results show that the Ct binds to lipid vesicles in a Ca^2^_+_-dependent manner, with higher Ca^2^_+_ levels increasing binding efficiency.

Surface analysis reveals that Ca^2^_+_ triggers a conformational change not only to free the Nt and Ct domains but also to alter their tridimensional structure. We show, for the first time, that Ca^2^_+_ binding causes local rearrangement of the Ct, making the opposite side more neutral and positively charged, ready to bind membranes. Moreover, Ca^2^_+_ ions form a positive pattern ideal for RNA binding. Our results also suggest that Ca^2^_+_ induces Nt structural changes essential for RNA and membrane interactions. To our knowledge, this is the first evidence showing that Ca^2^_+_ binding causes structural changes in the Nt and Ct domains of annexins. These rearrangements are essential to expose regions needed for partner binding.

Several studies have shown annexins interact with lipid membranes via Ca^2^_+_ (7,25), with ANXA5(19) and ANXA2(26) binding phospholipids through Ca^2^_+_-dependent residues. These findings, however, involved individual phospholipids, not entire membranes. This led to the hypothesis that annexin’s convex side bound to Ca^2^_+_ interacts with lipids membranes. Our results suggest an alternative hypothesis, especially for ANXA11, where these conformational changes on the concave side are crucial for membrane binding, not RNA interaction. This is supported by the AlphaFold model of the Ct:RNA complex, which shows contacts mainly with Ca^2+^ on the convex side (Fig. S3).

Complementing the lipid-binding studies, our investigation revealed that the Nt and Ct also bind RNA in a Ca^2+^-dependent manner. The Ct exhibits a higher affinity for lipids and RNA whereas the Nt shows weaker affinity for these ligands, which may result from limited sequence specificity and the request of other protein partners, yet both interactions are abolished without Ca^2+^, underscoring a shared regulatory dependence from this ion. These bindings were already investigated mainly in different full length annexins(22) and by computational methods that suggested plausible RNA binding motifs within the first 50 amino acids of the Ct(27). On the other hand, information on the Nt affinity for RNA has just been analysed only *in vivo* study(3). We believe that our findings, which offer more detailed insights into these interactions and their regulation by Ca^2^_+_, are essential, combined with existing literature, to elucidate the molecular mechanisms through which the individual Nt and Ct contribute to ANXA11’s function.

To gather information necessary to elucidate further ANXA11 complex tethering function we demonstrated that both Nt and Ct can bind RNA and lipid membranes simultaneously in a Ca^2^_+_-dependent manner. This evidence highlights the need to identify two distinct surfaces where RNA and liposomes bind. Based on the domain surface rearrangement observed through MD simulations, we hypothesise that RNA binds to the convex Ca^2^_+_ binding surface, while membranes attach to the concave side.

The dynamic relationship between the Nt and Ct in binding to RNA and lipid membranes reveals a complex, yet coordinated, mechanism of molecular recognition that is heavily influenced by Ca^2+^ ions. Analyzing their respective affinities and the simultaneous binding sheds light on the multifaceted functions of ANXA11 within cellular processes. In Figure 8, we propose a sequence of events in which Ca^2^_+_ binds to the Ct, causing the release of Nt. Subsequently, due to their differing binding affinities, the Ct will associate with a lysosome and RNA. The Nt will then follow to assemble the complex, first binding with RNA and then with the lysosome that finally is directed toward the molecular motors. This intricate hierarchy, regulated by Ca^2+^, enables precise modulation of ANXA11’s binding to RNA and lysosomes. Such multi-layered regulation facilitates the formation of a large complex with an extensive interaction surface. This is an essential requirement to resist hindrance encountered while transported.

**Figure 8.**
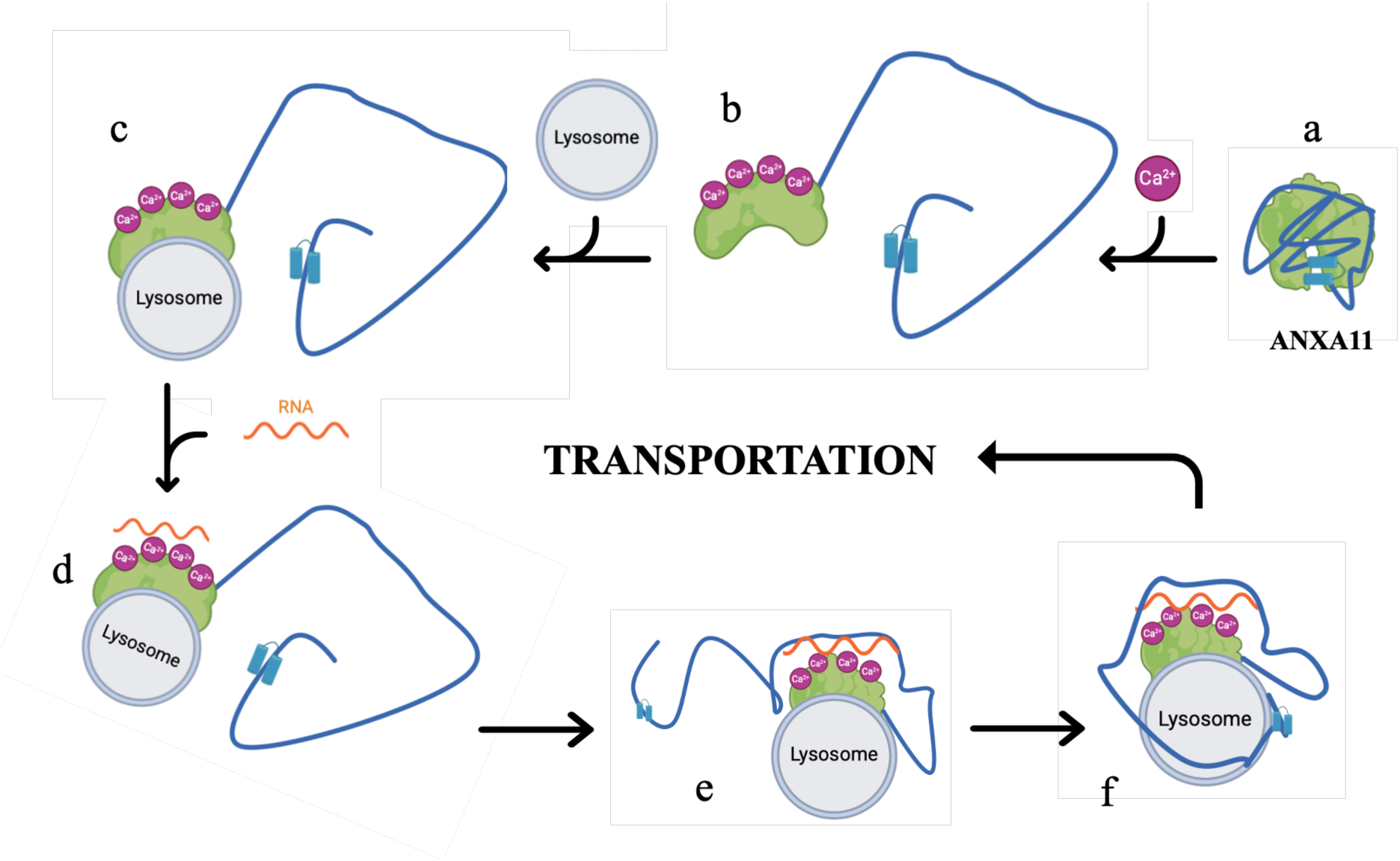
Cartoon depicting sequential events in the formation of the ANXA11 complex with RNA and lysosome. **a**, Initially, ANXA11 is in a closed conformation (Ct domain is in green colour, Nt domain in blue colour). **b**, Upon interaction with Ca^2+^, it undergoes a conformational change, adopting an open state. **c**, This facilitates increased binding affinity of the Ct domain to the lysosome. **d**, The Ct domain also shows increased affinity for RNA. **e**, The Nt domain binds to RNA as well. **f**, In addition, the Nt domain interacts with the lysosome. **g**, The resulting stable and expanded complex is then directed towards the molecular motor for transportation.

In conclusion, dissecting ANXA11 into its constituent domains has highlighted the regulatory mechanisms governing its function. The calcium-dependent modulation of intra- and intermolecular interactions revealed a sophisticated control system that likely responds dynamically to cellular Ca^2+^ signals, thereby regulating RNA transport and organelle tethering. These insights not only deepen our understanding of ANXA11’s molecular behavior but also have implications for pathological conditions like ALS, where dysregulation of these mechanisms may contribute to disease pathology. Continued structural and functional dissection at the domain level promises to uncover further details necessary for targeted therapeutic strategies.

## Material and Methods

### ANXA11 Nt, Ct and Nìdmg domain production

Variations of previously published purification protocols were used to obtain soluble, highly pure isolated ANXA11 Ct (residues 201–505) (28) and Nt (residues 1–191) (11) domains, while a novel protocol was adopted for the Nt_D40G_ mutant (p.D40G mutant, residues 1–191). The N-terminal (Nt, residues 1–191), C-terminal (Ct, residues 201–505), and Nt_D40G_ domains of ANXA11 were produced as thioredoxin fusion proteins, a strategy employed to enhance protein solubility, using constructs cloned into a pET His6 TEV LIC plasmid. Expression was carried out in *E. coli* BL21 (DE3) cells grown in LB supplemented with 100 μg/mL ampicillin at 37°C until OD600 reached 0.6, followed by induction with 1 mM IPTG at 20°C overnight. Cells were lysed in buffer containing 20 mM Tris-HCl pH 7.5, 150 mM NaCl for Ct or 20 mM NaCl for Nt and Nt_D40G_, 10 mM imidazole, 1 mM DTT, and 1 mM PMSF. Soluble proteins were purified by Ni-NTA chromatography using an elution buffer matching the lysis buffer but containing 300 mM imidazole. When necessary, the thioredoxin tag was removed by TEV protease digestion overnight at 4°C during dialysis against SEC buffer (20 mM Tris-HCl pH 7.5, 150 mM NaCl for Ct, and 20 mM NaCl for Nt and NtD40G). Proteins were further purified by reverse Ni-NTA chromatography and size-exclusion chromatography on a Sephadex 75 10/300 column. Chromatograms showed a single peak corresponding to the monomeric form of each domain.

### SDS-PAGE

Protein samples were diluted at a 1:1 ratio with denaturing buffer (0.125 M Tris-HCl, pH 6.8, 50% glycerol (v/v), 17 g/L SDS, 0.1 g/L bromophenol blue) in the presence of 2- mercaptoethanol and heated at 100 °C for 5 min. Electrophoretic runs were carried out in a MiniProtean Tetra Cell apparatus (Bio-Rad, Milan, Italy) on a 15% acrylamide-bisacrylamide (37.5:1 ratio) gel, using Tris/glycine buffer system at pH 8.3. Gels were stained with GelCode™ Blue Safe Protein Stain (Thermo Fisher Scientific, Milan, Italy).

### Western blot Analysis

Protein samples were separated by SDS-PAGE and transferred onto a PVDF membrane (Immobilon-P, Millipore, USA) using Trans-Blot Turbo (Bio-Rad, USA). Blocking was performed in 5% non-fat dry milk in TBS-T. The membrane was incubated overnight at 4°C with Rabbit Anti-His Tag antibody (1:1000, Thermo Fisher Scientific, USA) in 5% BSA/TBS- T. After washing, it was incubated for 2 hours at room temperature with HRP-conjugated Goat Anti-Rabbit IgG (1:10000, Thermo Fisher Scientific, USA) in 5% non-fat dry milk/TBS-T. Protein bands were detected using Clarity Western ECL Substrate (Bio-Rad, USA) and imaged with the ChemiDoc Imaging System (Bio-Rad, USA).

### Circular Dichroism (CD)

CD spectra were acquired using a Jasco J-815 spectropolarimeter equipped with a Jasco PFD- 425S Peltier system. Measurements were performed in a 0.1 mm path length quartz cuvette under a continuous nitrogen 5.5 flow to minimize oxygen interference at wavelengths near 200 nm. Protein samples were prepared at 0.1 mg/mL in 20 mM Hepes buffer, pH 7.5. Spectra were recorded in the 190–260 nm range at a scanning speed of 100 nm/min. Each spectrum represents the average of three consecutive scans. Secondary structure estimation was performed using K2D2 and K2D3 algorithms. Thermal unfolding profiles were recorded by monitoring the CD signal at 222 nm from 5°C to 75°C at a heating rate of 1°C/min. Spectra were collected at 5°C, 25°C, 30°C, 40°C, 50°C, 60°C, and 75°C.

### Differential Scanning Calorimetry

The DSC experiments were performed using VP-Capillary DSC, Malvern Instruments Ltd. Worcestershire, UK. The samples were placed in the calorimeter in a 200-μl sample cell against a 200-μl -ml reference cell that was filled with a blank solution consisting of the Ct of protein containing its own buffer. Both tdx-Ct fused protein and thioredoxin were used at 14 µM in all experiments. The cells were equilibrated inside the calorimeter for 10 min at 25 °C before heating up to the final temperature at a given rate. Cycles of cooling and reheating of the samples were performed to obtain the background for buffer subtraction or to test the hypothetical refolding. Replicate runs did not vary more than 0.25 °C. The denaturation temperature T_m_ and the ΔH were determined by fitting the data with the ORIGIN embedded program suite. To perform the analysis at different scan rates, the cells were pre-equilibrated with at least 10 cycles of buffer in the same setup of the experiment.

### Fluorescence Measurements of the Ct with Ca^2+^

Fluorescence measurements were performed on a PerkinElmer® LS-55 spectrofluorimeter equipped with a liquid thermostatted cell holder set at 25 °C. Excitation was set at 277 nm and emission spectra were recorded from 292–400 nm. Protein concentration was 6 µM, scan speed 50nm/min, with excitation/emission slit widths of 5.0 and 8.0 nm. Protein samples were prepared in 200 µL of buffer (20 mM HEPES, 150 mM NaCl, pH 7.5). Calcium titrations were carried out by successive additions of CaCE to reach final concentrations from 0 to 32 µM, without altering protein concentration, and the intrinsic fluorescence of tyrosine residues in the protein structure was used to monitor changes in the microenvironment upon calcium binding(29,30) and each addition, solutions were mixed and spectra acquired. Control experiments were performed by adding equivalent buffer volumes instead of CaCk

### Liposomes Preparation

Liposomes were prepared as *in vitro* models of biological membranes by thin lipid–hydration method mixing together L-α-phosphatidylcholine (Soy PC, Merck), 1,2-dimyristoyl-*sn*- glycero-3-phosphoethanolamine (DMPE, Merck) and cholesterol (CHOL, Merck) in 85:10:5 molar ratio. Lipids were dissolved in chloroform and evaporated by a rotary evaporator. The resulting lipid film was dried further under vacuum for 1 h to ensure complete solvent removal and then hydrated with 4-(2-hydroxyethyl) piperazine-1-ethanesulfonic acid (HEPES) buffer (20 mM, pH 7.5), vortexed and bath sonicated. The hydrated lipid suspension was sequentially extruded (Extruder, Lipex, Vancouver, Canada) at 30°C under nitrogen through different size polycarbonate filters (Costar, Corning Incorporated, NY), forming liposomes of approximately 600 nm (LipoA) and 150 nm (LipoB) in diameter. The liposome size and size distribution were determined by dynamic light scattering (DLS). The phospholipid amount was determined in each liposomal preparation by phosphate assay after destruction with perchloric acid(31). All liposomes were stored at 4 °C.

### p-NPL micelles preparation

The micelles were prepared by dissolving 16 mg of p-NPL (*Sigma-Aldrich^®^*) in 2 mL of anhydrous isopropanol, then diluting 0.25 mL of this stock solution in 4.75 mL of buffer (20 mM Tris, 150 mM NaCl, 0.5% v/v Triton X-100, pH 7.5) preheated to 50°C to reach a final p- NPL concentration of 1.24 mM.

### Pull-down Assay

For the liposome binding, 100 µg of protein were incubated with 100 µL of liposomes (approximate concentration of 10 mM) at 4°C for 30 minutes. The liposomes were prepared in-house, as described in the Materials and Methods section. After centrifugation (7000 rpm, 30 min, 4°C), the pellet was washed with Ct buffer (20 mM Tris-HCl, 150 mM NaCl, pH 7.5) or Nt buffer (20 mM Tris-HCl, 20 mM NaCl, pH 7.5) until the Bradford assay showed no further color change. The pellet was then resuspended in 30 µL of 2X Laemmli Sample Buffer, centrifuged (14,000 rpm, 5 min), and denatured (100°C, 10 min).

For the Poly-U Acid Agarose resin (Sigma-Aldrich®) binding, 25 µL of resin was incubated with 100 µg of protein at 4°C for 30 minutes. After centrifugation (1000 rpm, 1 min), the resin was washed with Ct or Nt buffers until the Bradford assay showed no further color change. The pellet was resuspended in 25 µL of 2X Laemmli Sample Buffer, centrifuged (15,000 rpm, 5 min), and denatured (100°C, 5 min).

For the Ni-NTA resin pulldown, 25 µL of Ni-NTA resin (Qiagen®) were incubated with protein at 4°C for 30 minutes. After centrifugation (1000 rpm, 1 min), the resin-bound protein was incubated with 10-mer Poly-U RNA labeled at the 3’ end with 6-FAM at 4°C for 30 minutes. Following a single wash with Ct or Nt buffer, the resin was centrifuged (1000 rpm, 1 min). The bound protein was then eluted from the Ni-NTA resin by incubation with 500 mM imidazole, followed by centrifugation (1000 rpm, 1 min). The fluorescence of the supernatant (excitation at 490 nm, emission at 520 nm) was measured to assess RNA binding. The experiment was performed in triplicate.

For the liposome-RNA three-component assay, liposomes with a mean diameter of ∼600 nm were prepared following the same protocol described above. After protein adsorption (Ct or Nt) on the liposomes, 10-mer Poly-U RNA labeled at the 3’ end with 6-FAM was added and incubated at 4°C for 30 minutes. Samples were centrifuged (7000 rpm, 30 min, 4°C), and the pellet was washed with Nt or Ct domain buffers until no further protein was detectable by Bradford assay. The relative fluorescence of the pellet was then measured to detect the presence of bound labeled RNA using maximum excitation at 490 nm and emission at 520 nm. The experiment was performed in triplicate.

For the three-component interaction assay, 25 µL of Poly-U Acid Agarose resin (Sigma-Aldrich® Poly-U Acid Agarose) was incubated with 100 µg of Ct or Nt and para-nitro phenillaurate (p-NPL) at 4°C for 30 minutes. Samples were centrifuged (1000 rpm, 1 min, 4°C) to remove the supernatant. Subsequently, they were washed with Nt or Ct domain buffers several times until protein content was not detectable any further by Bradford assay. Finally, 20 µL of NaOH 6N was added to the samples to release p-nitrophenol. The sample was centrifuged (1,000 rpm, 1 min), and the absorbance was measured at 410 nm using a NanoDrop™ Colibri (Titertek-Berthold, Berthold Detection Systems GmbH, Germany) to measure the para-nitrophenol released. The experiment was performed in triplicate.

### Microscale Thermophoresis measurements

The binding affinity between the Ct and Nt domains, as well as the interactions of Ct or Nt with LipoB and RNA, and the interaction of the Nt_D40G_ variant with Ct, were investigated by microscale thermophoresis (MST) using a Monolith NT.115 instrument (NanoTemper Technologies, Munich, Germany). Measurements were performed using the Nano Red LED at 100% excitation power and MST power at 40%. Ct was labeled with the Protein Labeling Kit RED-NHS 2nd Generation (MO-L011, NanoTemper Technologies, Munich, Germany), yielding ∼7 µM labeled protein with a labeling degree of ∼0.6, while Nt was labeled similarly, yielding ∼3 µM labeled protein with a labeling degree of ∼0.34. Ct/Nt binding was measured using 5 nM labeled Ct and Nt concentrations ranging from 0.75 nM to 25 µM. Ct/NtD40G binding was measured under the same conditions as Ct/Nt. Ct/liposome binding was tested with 10 nM labeled Ct and LipoB concentrations ranging from 3 nM to 100 µM. Similarly, Nt/LipoB binding was measured with 10 nM labeled Nt and LipoB concentrations ranging from 1.8 mM to ∼55 nM. For RNA binding assays, Ct was labeled with the His-Tag Labeling Kit RED-tris-NTA 2nd Generation (MO-L018, NanoTemper Technologies, Munich, Germany), and binding was measured with 5 nM labeled protein and RNA poly-U 10-mer concentrations ranging from 30.5 nM to 62.5 µM; the same procedure was applied to Nt. All measurements were carried out at 20°C in 20 mM HEPES buffer, pH 7.5, with 0.05% Tween- 80, and data were analyzed using MO Affinity Analysis software v3.0.5 (NanoTemper Technologies, Munich, Germany).

### Isothermal titration calorimetry measurements

The calorimetric titrations were performed in a VP-ITC microcalorimeter (MicroCal, Malvern-Panalytical, Malvern, UK). A protein solution of 13.5 µM (Ct) or 10 µM (Nt) in the calorimetric cell was titrated with CaCl_2_ solutions previously prepared in chemically matched buffer (Tris-HCl 20 mM, NaCl 20 mM, pH 7.5). For the Ct domain, a 540 µM CaCl_2_ solution was used as titrant, whereas for the Nt domain a 100 µM CaCl_2_ solution was employed. A series of 4-µL injections were performed with a spacing of 150 s, stirring speed of 500 r.p.m., and reference power of 10 µcal·s^−1^. The heat evolved after each ligand injection was obtained from the integral of the calorimetric signal and normalized by the moles of protein injected. The heat due to the binding reaction was estimated as the difference between the observed heat of reaction and the corresponding heat of dilution by including an adjustable term in the fitting routine accounting for the background injection heat. The experimental data for the Cterm were fitted using a model with two classes of independent binding sites (n = 2 per class), reflecting the biphasic nature of the isotherm, whereas the Nt displayed a typical sigmoidal binding curve that was best described by a one-site binding model (n = 1). In both cases, the fitting yielded the association constant (K_a_, with K_d_ = 1/K_a_), the interaction enthalpy (ΔH), and the apparent stoichiometry (n).

### Field Emission Scanning Electron Microscopy (FE-SEM) combined with Energy Dispersive X-ray Spectroscopy (EDS) analysis

FE-SEM measurements have been carried out by using a Tescan S9000G FESEM 3010 microscope (Tescan Orsay Holding a. s., Brno- Kohoutovice, Czech Republic) working at 30 kV, equipped with a high brightness Schottky emitter and with EDS probe fitted with an Ultim Max Silicon Drift Detector (SDD, Oxford, UK). For the analyses, one drop of each prepared sample was deposited onto an aluminium stub coated with a conducting adhesive and subsequently left to dry in the air at room temperature. To avoid charging effects, the samples were then metallized with Cr (ca. 5 nm) using an Emitech K575X sputter coater (Quorumtech, Laughton, East Sussex, UK) and further inserted into the chamber by a motorized procedure. Images were taken by acquiring either secondary electrons (SE) or backscattered electrons (BSE) working at 5 KeV to avoid modification or damage of the sample during the observation and adopting the in-beam high resolution mode. Representative FE-SEM images and corresponding EDS spectra were collected in different regions of each sample. By this approach, morphology and elemental composition of the samples made up by ANXA11 Ct (20 µM) + lipid vesicles 200 µM and ANXA11 Nt (20 µM) + lipid vesicles (200 µM) were deeply investigated. The effect of Ca^2+^ was studied by adding a CaCl_2_ solution (400 µM) to both samples.

### Computational model predictions

In the absence of an experimental ANXA11 (UniProt entry P50995) structure, a query on the AlphaFold Protein Structure DB built on AlphaFold 2 (available at https://alphafold.ebi.ac.uk)(32,33) was performed. Then, taking advantage of the recently updated version of the AlphaFold AI model (23) — available as a web server at https://alphafoldserver.com/about — we also predicted ANXA11 structure in the presence of Calcium ions.

By means of the AlphaFold Server different predictions of ANXA11 (from Ile37 to Asp505) were attempted, in the presence of four Ca^2+^ ions and in their absence. Each job (producing five different predicted models) was repeated three times on the platform using a different randomly generated seed, for each chosen molecule combination. This allowed us to make a first comparison between the prediction in the absence of Ca^2+^ ions generated by AlphaFold 2 (obtained from the DB with entry P50995, Fig. S2a), and the predictions by Alphafold 3. Careful visual investigation of the models was performed to verify the quality of the predictions (e.g. absence of steric clashes, loop conformations, calcium ion coordination). The resulting structures with and without Ca^2+^ ions (Fig. S2b and S2c respectively), represented the best obtained models in terms of predicted template modeling score and interface predicted template modeling score, and led to a different outcome in terms of the occurrence of the quaternary interactions between the Nt and the Ct domain.

In order to shed light on the role of Calcium in ruling the interaction between the Ct domain and the HtH motif of the N-terminal domain), the following models were built and simulated: *Nt-Ct, ANXA11 in the absence of Calcium ions (removing the amino acids from 67 to 190, in order to focus only on the interaction between the HtH motif and the Ct domain); *Nt-Ct- Ca^2+^, where 4 Calcium ions were added also grafting the so-called AB-loops, coordinating the ions – i.e. Lys214-Asp219 (Ca1, Fig. S2f), Lys286-Asp291 (Ca2, Fig. S2g), Gly368-Asp375 (Ca3, Fig. S2h), Arg445-Lys450 (Ca4, Fig. S2i) – accordingly; Ct-Ca^2+^, the control model of the Ct domain in the presence of Calcium ions, but in the absence of the Nt HtH motif. Moreover, taking the resulting *Nt-Ct model and removing the side chain of the Asp40, easily replaced by a H atom, the disease-associated p.D40G variant(12) was obtained, thus carrying a glycine in place of an aspartic acid residue at position 40 of the HtH motif in the N-terminal domain, thus named *Nt_D4øG_-Ct. An additional AlphaFold 3 prediction was tried out combining ANXA11, four Ca^2+^ ions and two RNA poly-U 10-mer strands (Fig. S3).

### Molecular dynamics (MD) simulations

The selected predicted protein models were let explore the conformational space available to them in simulations (via equilibrium MD), with the GROMACS 2024 engine (34); a widespread protocol(35,36) (details in SI) was applied to set up, minimize and equilibrate the systems before the 2.5 µs-long production simulations (2 replicas for each investigated system) using the amber14sb force field(37), the velocity-rescaling thermostat(38) and the stochastic cell rescaling barostat(39).

For the analysis of the trajectories we employed multiple tools: the Gromacs suite (40) was used for the RMSD, RMSF, contact analysis and the cluster analysis; the Adaptive Poisson-Boltzmann Solver(41) plugin included in PyMol (v 2.5.2) was used to perform the electrostatic analysis (with default parameters); the Matrix of Low Coupling Energies method (42) was employed for the so-called epitope mapping, in the same fashion as in Casale et al.(43). All the details are provided in SI.

## Author contributions

S.A. conceived and supervised the project. A.F., G.D.N. and L.A. carried out the Pull-down assays, the MST measurement, electro-microscopy and Isothermal calorimetry. P.O. and E.R. established the protein purification protocols. V.B. and S.Ar. were involved in lipids vesicles preparation. G.L.C. and G.F.G. produced the DLS and fluorescent measurement. M.Mz. guided in carrying out the electro-microscopy measurements. A.V.C. helped with the data analysis of the ITC. A.D.S. helped analyse the RNA binding data and with the manuscript preparation. F.D.P. carried out the molecular dynamics and with the manuscript preparation. R.S. performed bioinformatic analyses. F.S. guided both bioinformatic analysis and molecular dynamics. M.M. and F.D.P. carried out the secondary structure characterization. S.O.B and F.P. verified the analytical methods. All authors provided critical feedback and helped shape the research, analysis and manuscript.

## Supporting information

Zip file with supplementary information

