## Supplementary material for "Dissecting Annexin-A11 into its functional domains revealed calcium as a key regulator for RNA transport and its association with ALS": Zip file with supplementary information: supplementary_material .pdf

### DSC spectra of Thioredoxin alone

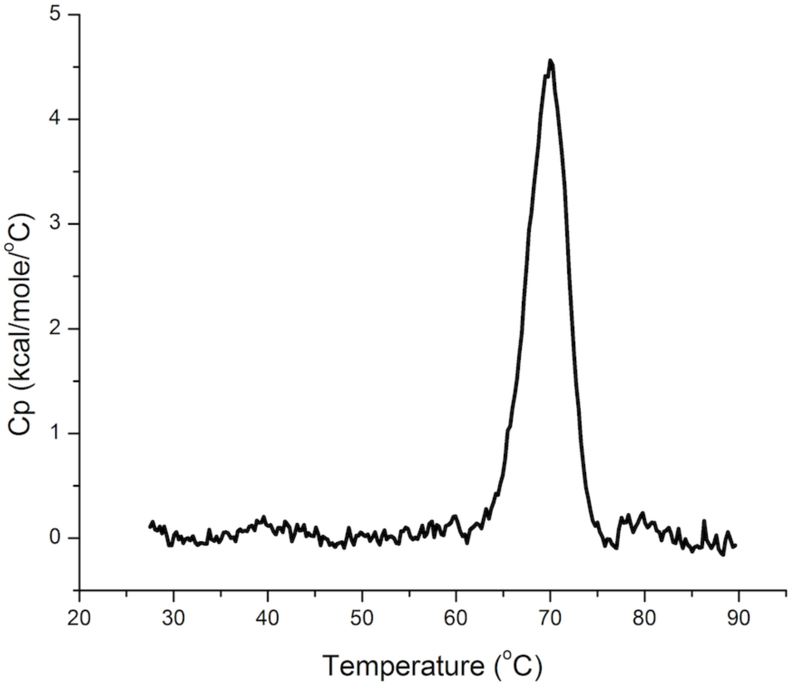


**Fig. S1. DSC thermogram of Thioredoxin in the absence (black) and presence (red) of $Ca^{2+}$.** The first derivative highlights the thermal transition midpoint around 70 °C

### ANXA11 N-terminal and C-terminal domain prediction and characterization

Taking the structural prediction deposited in the AlphaFold Protein Structure Database (DB) as reference model for ANXA11 (given that it can be considered the best possible prediction generated by Alphafold 2, Fig. S2a), we compared the new structures resulting from the prediction jobs performed via the AlphaFold 3 server, both in the presence and in the absence of Calcium ions, with the former one; it is worth to notice that at variance with the previous version of the generator, the AlphaFold 3 deep-learning framework allows you to model a structure consisting of many biological molecules, hetero-complexes, including proteins, nucleic acids and ions(44). Thus, this *in silico* investigation started from the analysis of the AlphaFold 3 resulting models: the main and clear difference between the ANXA11 predictions with 4 or no $Ca^{2+}$ was in the tertiary interaction between the Helix-turn-Helix (HtH) motif of the N-terminal (Nt) domain and the C-terminal (Ct) one: present in the absence of the ions (Fig. S2b) (as in the entry at the AlphaFold DB, https://alphafold.ebi.ac.uk/entry/P50995, Fig. S2a), but completely depleted in the presence of them (in 15 out of 15 predictions, Fig. S2c).

To better characterize the calcium-binding feature of the ANXA11 Ct domain, we performed a comprehensive sequence (Fig. S2d) and structural alignment of the first two Ct Calcium binding repeats (Fig. S2e - repeats 1 and 2 are colored in green) versus the last two repeats (Fig. S2e - repeats 3 and 4 are colored in yellow). Our analyses revealed a high degree of sequence homology and structural similarity between these two sets of repeats (Fig. S2d and S2e, respectively) that almost perfectly align two by two.

Indeed, structural homology suggests an evolutionary pattern where the Ct might have been composed of a tandem repeat of two units. Subsequent dimerization events may have facilitated the emergence of a four-repeat domain, thus enhancing calcium affinity and functional complexity.

A detailed comparison of the calcium-binding loops revealed notable sequence-specific differences between the four ANXA11 calcium-binding sites (Fig. S2). At variance with ANXA4, exhibiting conserved sequences of all four loops(45), ANXA11 displays slightly distinct loop sequences (Fig.  S2d), which may concur to a finer calcium regulation.

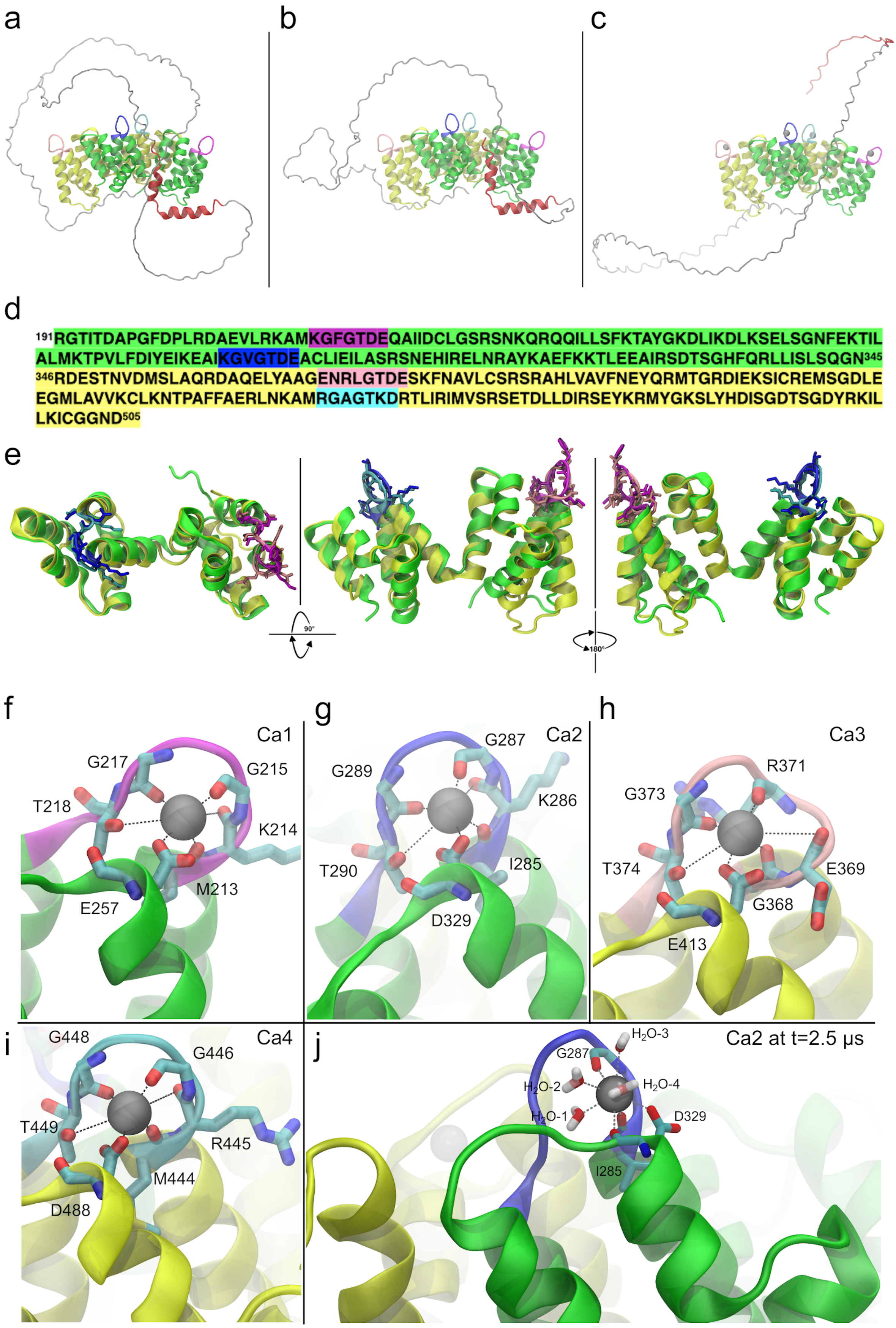


**Figure S2.** ***In silico*** **characterization of the ANXA11 N-terminal and C-terminal domains. a**, AlphaFold 2

ANXA11 full-length prediction as deposited in AlphaFold DB (entry: P50995), the HtH motif of the Nt domain is colored in red, repeat 1 and 2 in green and repeat 3 and 4 in yellow. **b**, AlphaFold 3 ANXA11 full-length prediction in the absence of Calcium. **c**, AlphaFold 3 ANXA11 full-length prediction in the presence of four $Ca^{2+}$ ions (as silver spheres). Sequence (**d**) and structural (**e**) alignments of the dimer of ANXA11 Ct repeats 1 and 2 (green) versus 3 and 4 (yellow). The AB loops, corresponding to the calcium binding sites of each repeat, are highlighted in different colors (magenta, blue, pink and cyan for the repeats 1, 2, 3 and 4 respectively). The focus on each $Ca^{2+}$ binding site is also shown (panels **f**, **g**, **h**, **i**) with the labelled residues coordinating the ions, together with an example of the typical Calcium coordination at the end (t=2.5 µs) of the Ct-$Ca^{2+}$ Replica1 simulation (**j**).

Another important feature emerging from the predictions in the absence of Calcium was the folding of the HtH motif: in 12/15, the prediction led to an effective HtH structure (in perfect agreement with the previously mentioned AlphaFold 2 prediction deposited in the DB), whereas in the remaining 5 predictions the αα from I37 to G66 were folded in a unique helix (3/15) still partially interacting with the Ct domain at the same spot.

At last our attention was drawn to the amino acids coordinating the 4 $Ca^{2+}$; in a near-future, solving the structure of the ANXA11 in complex with Calcium, it would be highly probable to spot those ions in contact with the already mentioned loops because, in good accordance among the 15 generated models, the ions were placed at the same sites near the already mentioned loops, i.e. [K214-E220 (magenta, Figure S2f), K286-E292 (blue, Fig. S2g), E369-E376 (pink, Fig. S2h), R445-D451 (cyan. Figure S2i)].

For the sake of completeness, we also included the last conformation of the calcium-binding loops of the repeat 2 sampled at the end of the Replica1 Ct-$Ca^{2+}$ control trajectory (Fig. S2j), as an illustrative example of the Calcium ion coordination in the presence of water after 2.5 µs of MD simulation (the conformation of the other three $Ca^{2+}$ binding sites were very similar to the one shown at Fig. S2j, similar also to the ones resulting from the *Nt-Ct-$Ca^{2+}$ simulations). The presence of three backbone carbonyl oxygens from the AB loop residues, the side chain of an acidic amino acid (D329 in this specific case) downstream from the DE loop and (at least) two water molecules (4 in total in this case), coordinating the Calcium ion, allowed us to recognize it, as a canonical type II Calcium binding-site(46,47,48), like the other three sites on the other repeats.

Further, using AlphaFold 3, we predicted the full-length ANXA11 in complex with four $Ca^{2+}$ ions and two RNA poly-U 10-mer, in a similar fashion to our RNA-binding experiments (see main text). The resulting prediction, revealing the RNA strands in a tight interaction with the Ca-binding side of the Ct, is shown in Figure S3.

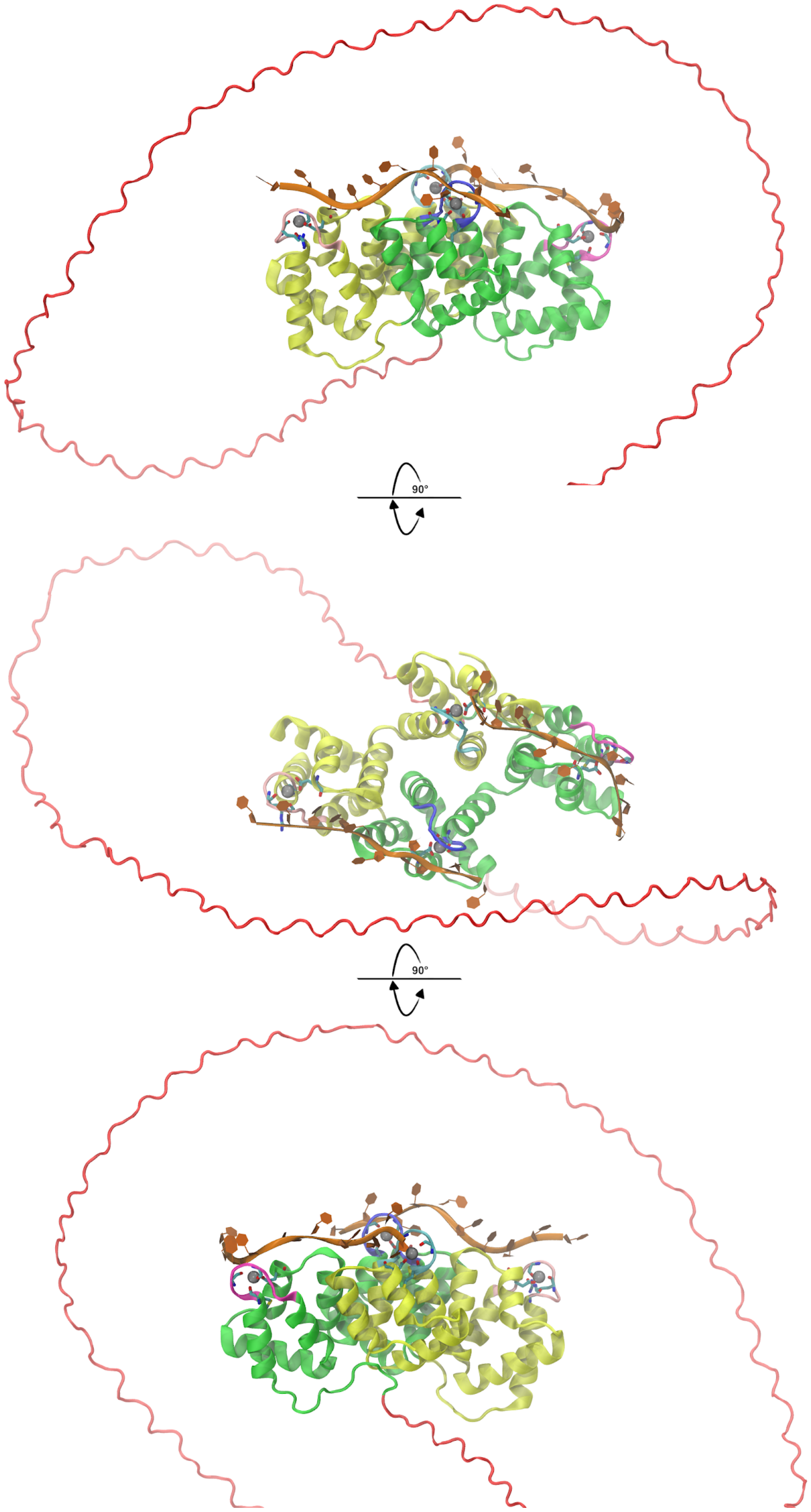

**Figure S3**. ***In silico* prediction of the full-length ANXA11 in complex with four Calcium ions and two RNA poly-U 10-mer by means of AlphaFold 3.** Three alternative views of the same model are shown, top view (mid panel) and side views (horizontal +/– 90° rotation, top and bottom panel respectively). The ANXA11 Nt was predicted as disordered, including the HtH in red, and not interacting with the Ct; the Ct repeats 1 and 2 are colored in green, repeats 3 and 4 in yellow, the AB loops, corresponding to the calcium (as silver spheres) binding sites are highlighted in different colors: magenta, blue, pink and cyan for the repeats 1, 2, 3 and 4 respectively; the two RNA poly-U 10-mer (in orange backbone and ribonucleotides) were predicted in tight interaction with both the $Ca^{2+}$ ions and the AB (and CD) loops: specifically, one poly-U interacts with repeat 1 and 4, and the other interacts with repeat 2 and 3.

### MD simulations protocol and analysis details.

In order to set up the systems for the MD simulations, the first choice concerned the force field: the amber14sb(49) was used to model the all-atom system; an octahedron-shaped box was built setting its length on the basis of the molecule longest axis, further adding an extra space of 1.2 nm between the solute and the box edges; the box was then solvated using the spc/e explicit model for water(50). The systems were completed upon addition of a saline solution (0.15 M KCl) and, eventually of the ionic concentration required to reach electro-neutrality ($K^+$), to reproduce the experimental conditions. First a minimization was carried out with the steepest descent algorithm and terminated as the maximum force decreased to less than $100\ \frac{kj/mol}{nm}$ to allow a protein global relaxation. The system equilibration was run afterwards to thermalize and pressurize the system: two the performed steps to this aim: 1) 1 ns NVT, in which the first 300 ps were employed to perform a simulated annealing to bring the temperature from 0 to 300 K, using the velocity-rescaling thermostat(51) with a 2 fs time step and keeping the complex restrained $[1000\ \frac{kj/mol}{nm}]$ in order to let the water properly arrange around it; 2) 4 ns NPT at 1.0 bar using the stochastic cell rescaling algorithm to keep pressure constant(52), with the second half used to gradually release the restraints on the complex heavy atoms in order to reach an unrestrained condition. The production simulation time of one of the two replicas of the system in the presence of the HtH Nt domain and in the absence of the calcium ions (*Nt-Ct) was doubled, till 5 $\mu s$, for testing purposes: in this condition the interaction between the HtH motif and the Ct domain was still lasting (data not shown), at variance with the *Nt-Ct-$Ca^{2+}$ simulations (see Results).

VMD 1.9.4(53) was used to visualize the trajectories and image rendering, but also to check the key interactions between the protein domains, and to monitor the distance between the calcium ions and their coordinating protein residues. The Gromacs suite(54) was used for the

RMSD, RMSF, contact analysis (the *hbond* module was employed setting a 3.5 Å threshold with the *-contact* flag in order to include all the putative electrostatic interactions), and the cluster analysis (default parameters). For both the deviation and fluctuation analysis, RMSD and RMSF were calculated over the entire simulation time. Both the RMSD and the cluster analysis were performed over a unique concatenated trajectory of the 2 independent replicas for each system; the RMSF instead was run over the independent trajectories. At last, on the structure representing the most populated cluster for each system (i.e. centroid), we also performed the electrostatic calculation and the putative epitope mapping. For the systems sampling the disruption of the HtH motif/Ct domain interaction, we splitted the trajectories in two parts and we made the cluster analysis on the bound portion and the unbound one separately.

The electrostatic analysis was performed taking advantage of the Adaptive Poisson-Boltzmann Solver(55) plugin included in PyMol (v 2.5.2) with default parameters, allowing us to display the results of the calculations as an electrostatic potential molecular surface; the epitope search, instead, aims at the identification of 3D substructure of interest that are prompt to interact with binding partners. These protein regions might be formed by distant segments of the primary sequence that come together in the folded protein, namely a conformational epitope. Most epitope prediction approaches rely on static structure derived from crystallographic methods, whereas the here employed Matrix of Low Coupling Energies (MLCE) method(56) takes advantage of the MD simulation ability to provide insight into protein motion overtime, capturing important interactions and conformational changes that may influence the recognition processes; the epitope mapping was performed here in the same fashion as in Casale et al.(57).

**MD simulation final conformations.**

Here we show the last conformation sampled in the four *in silico* systems (*Nt-Ct, *Nt-Ct-$Ca^{2+}$, Ct-$Ca^{2+}$, *$Nt_{D40G}$-Ct) at the end of the trajectories; this is useful to visualize the effect on the C-terminal domain of the presence/absence of both Calcium ions and the HtH motif of the Nt domain. In the *Nt-Ct system, at t=2.5 μs, in the absence of $Ca^{2+}$ the HtH is still attached to the Ct domain, in its initial position and almost in the initial conformation too (Fig. S4b, Video 2A). On the contrary, at the end of both the simulated replicas in the presence of $Ca^{2+}$, the unbinding process was completed and the final *Nt-Ct-$Ca^{2+}$ conformation showed the HtH motif no longer interacting with the Ct domain (Fig. S4a). The destabilization induced by calcium is crystal clear as the trajectories revealed the HtH domain fully dissociating from the Ct domain (Videos 1A and 1B).

The final conformation of the Ct-$Ca^{2+}$ (Replica1, Fig. S4c), control system, shows the Ct domain competence in keeping the $Ca^{2+}$ ions bound, thus decoupling the stabilizing effect of the Calcium from the destabilizing effect of the HtH motif, in the simulations (Videos 3A and 3B).

Moreover, in order to verify the effect of the D40G mutation in the HtH motif, the final conformation of the *$Nt_{D40G}$-Ct is also provided (Fig. S4d). As it resulted from the MD simulations, the mutation at position 40 possibly caused the unbinding of the HtH motif from the Ct domain in the absence of $Ca^{2+}$ ions (Videos 4A and 4B), thus bypassing the calcium regulation.

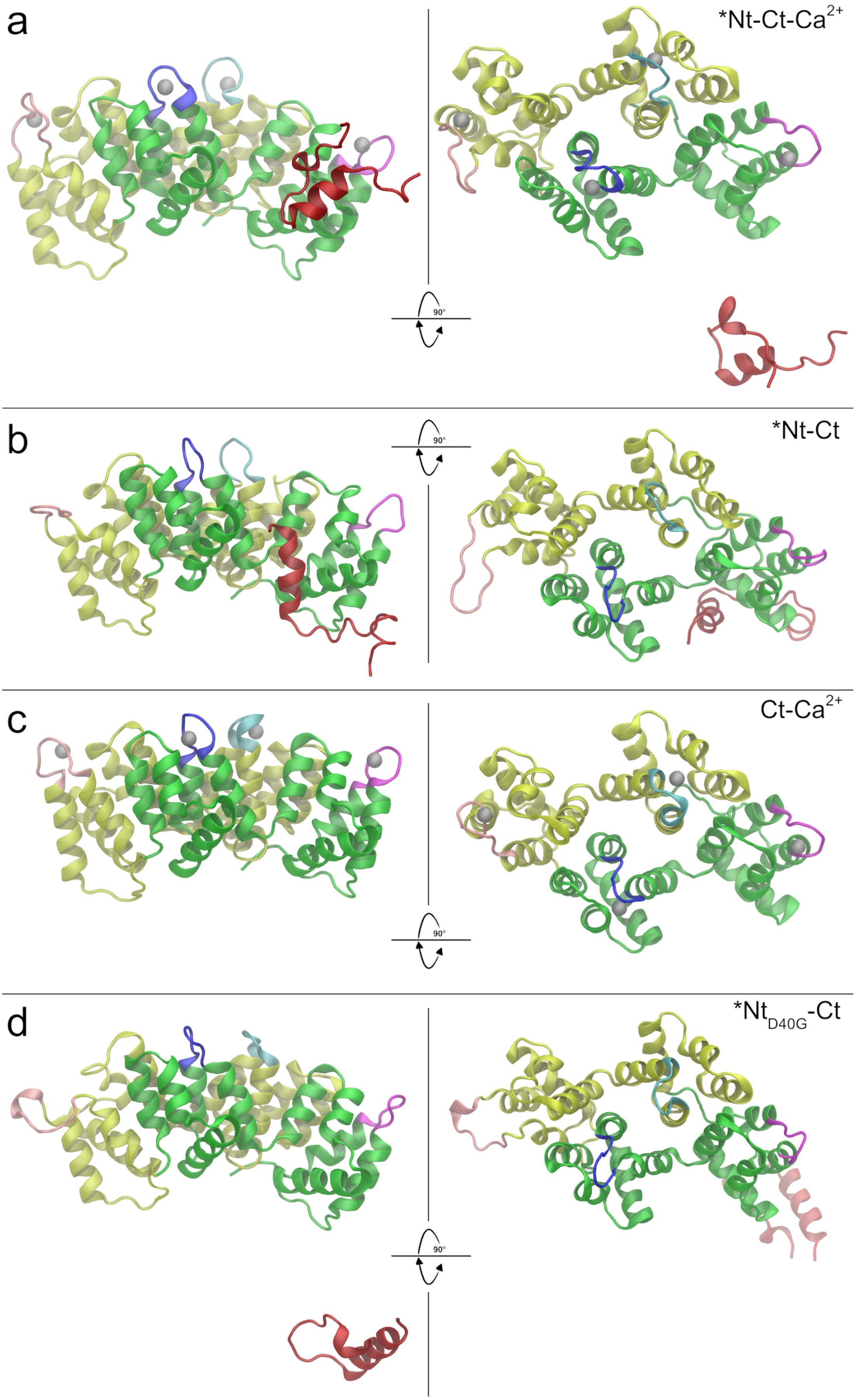
a
*Nt-Ct-Ca2+
90°
b
*Nt-Ct
90°
c
Ct-Ca2+
90°
d
*NtD40G-Ct
90°

**Figure S4. Final conformation at 2.5 $\mu$s for each system. a**, *Nt-Ct-$Ca^{2+}$, with the HtH motif not bound to the C-terminal domain. **b**, *Nt-Ct, with the HtH motif still bound to the C-terminal domain. **c**, Ct-$Ca^{2+}$, in the absence of the HtH motif and in the presence of $Ca^{2+}$ ions. **d**, *$Nt_{D40G}$-Ct, with the mutated HtH motif no longer interacting with the C-terminal domain also in the absence of Calcium. For completeness, Replica2 final conformation for each system is viewable at the end of the corresponding attached Videos 1B, 2B, 3B, 4B.

### Charge Density Analysis and Epitope Mapping

Given the surface charge density of the C-terminal on both the *Nt-Ct and the Ct-$Ca^{2+}$ systems presented in Figure 4a/c (main text), we performed the same analysis also on the *Nt-Ct-$Ca^{2+}$ (previously running the clustering on the C-terminal domain only on the portion of the two replicas downstream the HtH unbinding events): in the presence of the Calcium ions this system reported an intermediate behaviour between the *Nt-Ct and the Ct-$Ca^{2+}$; part of the exposed surface on the opposite side with respect to the $Ca^{2+}$ binding sites turned generally neutral, with few small partially negative spots still appearing on the surface (Fig. S5a/right); this was probably due to the short simulation time sampled after the HtH unbinding (after ~1.75 μs in Replica1 and after ~2.2 μs in Replica2) that was not sufficient to complete the conformational change, like in Ct-$Ca^{2+}$, thus not letting all the negative charged residues to be buried yet.

To some extent, the epitope mapping analysis by means of the MLCE method (13) might be complementary to the charge density analysis, revealing if a surficial charge density change (analyzing two different systems), corresponds to a different local propensity in having an interacting partner. As shown in Figure 4d, the Ct of the representative conformation resulting respectively from the *Nt-Ct and the Ct-$Ca^{2+}$ simulations presented a clear difference: A new epitope appeared (colored in red in Fig. 4d panel) exactly at the spot previously identified as the one in which the negative charges were buried in the presence of Calcium. On the contrary, the corresponding region in the absence of Calcium (Fig. 4b panel) was blank.

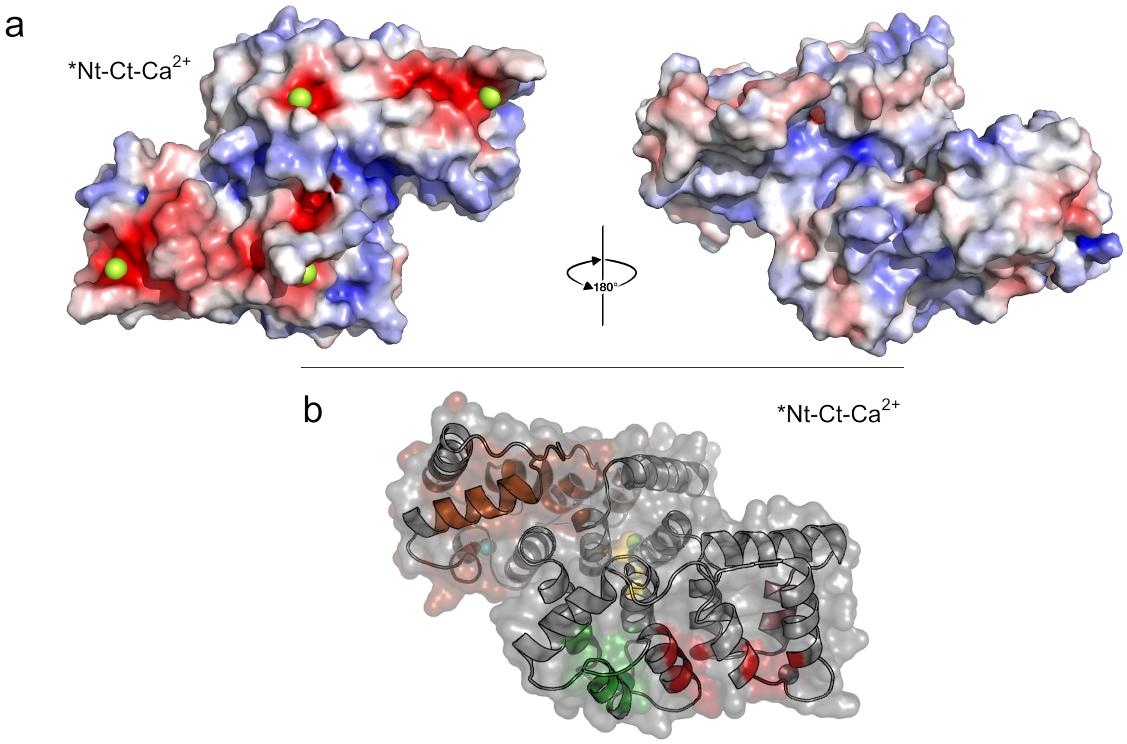


**Figure S5**. ***Nt-Ct-Ca$^{2+}$ charge density and epitope mapping. a**, Surface of the centroid structure of the most populated cluster resulting from the *Nt-Ct-Ca$^{2+}$ simulations (downstream of the HtH unbinding event) colored according to its charge density (red/negative, white/neutral, blue/positive); C-terminal domain is shown, on the left panel the surface with the four bound Ca$^{2+}$ (green spheres), the AB and CD loops exposed, on the right the opposite surface. **b**, Maps of the epitopes identified in the centroid structure of the *Nt-CtCa$^{2+}$ most populated cluster identified in the simulation downstream of the HtH unbinding event; the surface opposite to the Calcium binding sites is in the foreground, in red is colored an epitope located at a similar spot to the one appeared in the Ct-Ca$^{2+}$ map (Fig. 4d).
